# Co-reviewing and ghostwriting by early career researchers in the peer review of manuscripts

**DOI:** 10.1101/617373

**Authors:** Gary S. McDowell, John Knutsen, June Graham, Sarah K. Oelker, Rebeccah S. Lijek

## Abstract

The goal of this study is to shed light on the involvement of early career researchers (ECRs) during peer review of manuscripts for publication in journals. In particular, we sought to better understand how commonly ECRs contribute ideas and/or text to peer review reports when they are not the invited reviewer (“co-review”), and how commonly ECRs do not receive named credit to the journal editorial staff for these scholarly efforts (“ghostwrite”). First, we evaluated 1,952 publications in the peer-reviewed literature generated by exhaustive search terms that combined synonyms of “early career researcher” and “peer review” and found no previous studies about ECRs ghostwriting peer review reports. We then surveyed 498 researchers about their experiences with, and opinions about, co-reviewing and ghostwriting as ECRs. Three quarters of those surveyed have co-reviewed and most find it to be a beneficial (95% agree) and ethical (73% agree) form of training in peer review. Co-reviewing is the second most commonly reported form of training in peer review besides receiving reviews on one’s own papers. Half of survey respondents have ghostwritten a peer review report, despite the 4/5ths majority opinion that ghostwriting is unethical. Survey respondents report that the three major barriers to including co-reviewer names on peer review reports are: a lack of communication between PIs and ECRs; a false belief that co-authorship is for manuscripts but not peer review reports; and prohibitive journal policies that are out of alignment with current practice and opinions about best practice. We therefore propose recommendations for changing this status quo, to discourage unethical ghostwriting of peer review reports and encourage quality co-reviewing experiences as normal training in peer review.

## INTRODUCTION

Peer review of academic manuscripts is viewed as a fundamental scholarly activity to maintain the integrity of the scientific literature (Baldwin 2018; Tennant et al. 2017). Early career researchers (ECRs; Box 1: Definitions) often contribute to this peer review process. Indeed, in a recent survey shared on the INSIDE *eLife* blog that targeted ECRs in the life sciences, 92% of those surveyed reported undertaking reviewing activities (Inside eLIFE 2018). While it may be expected that ECRs review manuscripts jointly with or under the guidance of a principal investigator (PI; Box 1), more than half of survey respondents, including 37% of graduate students, reported reviewing a manuscript *without any assistance from their advisor.* Are these ECRs performing peer review independently as the invited reviewer? Or are they performing peer review on behalf of their advisor, the invited reviewer being named to the journal? In this case, is the journal editorial staff aware that someone other than the invited reviewer has contributed independently to the peer review report? If not, then these ECRs are participating in the practice of ghostwriting (Box 1). Combining the results of the INSIDE *eLife* survey with our own anecdotal observations of ECRs carrying out peer review without being identified to journal editorial staff, we came to the hypothesis that ghostwriting of peer review reports by ECRs is widespread.

### BOX 1: DEFINITIONS USED IN THIS STUDY

| BOX 1: DEFINITIONS USED IN THIS STUDY |  |
| --- | --- |
| **Early career researcher (ECR)** | We consider this to be anyone engaged in research who is not recognized as an independent leader of a research group, including: undergraduate, graduate, and postdoctoral researchers; junior research assistants, associates, and staff scientists. |
| **Principal Investigator (PI)** | Anyone recognized as an independent leader of a research group, including: professors, group leaders.<br>*Note:* We use this term to mean someone likely to be an invited reviewer due to their professional independence, including pre-tenure junior faculty (e.g. assistant professor in the US). We recognize that, in other contexts, pre-tenure faculty may also be categorized as ECRs. |
| **Co-reviewing** | Contributing ideas and/or text to a peer review report when one is not the invited reviewer. Equivalent to a co-author on a manuscript when one is not the corresponding author.<br>*Note:* We use this term to mean significant contributions to the peer review report, and so differentiate from casual or insignificant conversations about the manuscript under review that do not provide novel ideas and/or text to the peer review report. |
| **Ghostwriting** | Co-reviewing without named credit to the journal editorial staff.<br>*Note:* We use this term to mean <u>only</u> the identification of a co-reviewer to the journal staff in an identical manner to the identification and naming of the invited reviewers. We are not referring to the *public* naming of peer reviewers, or reviewers *signing* reviews, or other forms of open peer review which is beyond the scope of this study (Ross-Hellauer 2017). |

Indeed, co-reviewing and ghostwriting by ECRs appear to be understood as common practices in academia, summarized recently by Patterson and Schekman (2018):

> “It is common practice for busy group leaders to ask their more senior PhD students and postdoctoral fellows to help with peer review, but in too many cases these contributions go unacknowledged.”

This statement joins a growing discussion about co-reviewing and ghostwriting, which includes a workshop we led at the 2018 ASAPbio meeting on Transparency, Recognition, and Innovation in Peer Review in the Life Sciences (http://asapbio.org/peer-review/summary, (McDowell 2018)). However we wished to widen this conversation beyond those at attendance at a particular conference and facilitate an evidence-based dialogue about ghostwriting and, more broadly, about the participation of ECRs in peer review. We therefore sought to collect data that answers:

> How commonly do ECRs ghostwrite peer review reports, and what is their opinion of this practice? Do they find it valuable and ethical? Why do such practices take place? Are there any concerns that arise with this practice and could interventions be put in place to address these issues?

We performed a systematic review of the extant literature to determine if these questions had been previously addressed. Among the ~2000 publications in peer reviewed journals on the topic of “early career researchers” and “peer review”, we found no research articles reporting data specifically on peer review reports ghostwritten by ECRs. Given the lack of data currently available about this practice, its frequency, and the rationales behind it, we conducted a survey of peer review experiences and opinions that primarily targeted ECR communities in the biomedical sciences. **We describe here our findings from 498 survey respondents that give the first specific evidence of the frequency of peer review ghostwriting and the motivations behind it.** Survey data reveal logistical barriers that prevent ECRs from receiving credit for reviewing activities, such as a lack of clarity in journal policies concerning the expectations and reporting mechanisms for the participation of ECRs in peer review. Our data also suggest that overcoming these policy barriers would not be sufficient to fully involve ECRs in peer review, since there are also incorrect assumptions and cultural practices out of line with the values held by researchers about the involvement of ECRs in peer review. We therefore propose a series of policy and cultural changes supported by our data that would have the potential to ensure the inclusion, training, and recognition of ECRs’ scholarship in manuscript peer review.

## RESULTS: SYSTEMATIC LITERATURE REVIEW

We performed a systematic review of the peer-reviewed literature with the goal of identifying any previous studies on the role that ECRs play in the process of peer reviewing manuscripts. In particular, we wanted to know whether there was evidence in the literature of ECRs ghostwriting peer review reports. Since ghostwriting is, by definition, an outcome that results from a lack of documentation, transparency, and accountability, we hypothesized there would be little-to-no evidence-based literature on this topic. Investigating a null hypothesis is challenging and as a result we designed a comprehensive procedure based on evidence-based guidelines for systematic reviews using exhaustive search terms that combined any synonyms of “early career researcher” AND “peer review” (<u>prisma-statement.org</u>. (Moher et al. 2009), <u>Methods: Systematic Literature Review</u>).

Our search yielded 1,952 unique articles. Collected articles underwent two rounds of screening performed independently by 3 study authors using titles and abstracts to evaluate relevance to the topic of ECR co-reviewing and ghostwriting peer review reports (<u>Methods: Relevance screening; Appendix: Results of relevance screening for literature review</u>; Appendix Figure 1). 118 articles were considered relevant by at least one screener and 36 articles were considered relevant by two or more screeners. All 36 articles considered relevant by 2+ screeners then underwent a full text reading with specific attention being paid to: research question, motivation for article, method of study including details concerning study participants, relevant results and discussions, discussion of peer review and ECRs, and possible motivations for author bias. Of the articles uncovered by our search that were found not to be relevant to the topic of ECR involvement in the peer review of manuscripts, many discussed other forms of peer review outside the scope of publishing manuscripts (e.g. students in a classroom setting engaging in peer review of each other’s assignments as a pedagogical exercise).

### Lack of literature on ECR ghostwriting of peer review reports

We found 0 research articles in the peer-reviewed literature on the practice of ghostwriting of peer review reports by ECRs^1^. There was one publication, not a research article, that mentioned ghostwriting of peer review reports by ECRs: the announcement of peer reviewer training policies from the journal *eLife* that was quoted above and was published after the ASAPbio meeting mentioned above (Patterson and Schekman 2018). This policy announcement acknowledged the phenomenon of ECR ghostwriting and stated a journal policy that ECRs are eligible to act as peer reviewers of manuscripts submitted to *eLife.*

Of the remaining 35 articles that were considered relevant to the topic of ECR involvement in the peer review of manuscripts but *did not* address ghostwriting, many instead investigated the value of co-reviewing as a training exercise. We summarize the major themes from these articles in the <u>Appendix: Literature on ECR involvement in peer review as a training exercise</u>. None of these articles discussed the issue of named credit for scholarly labor, nor did they include information on the frequency of, or ECRs’ opinions about, ghostwriting in peer review.

## RESULTS: SURVEY OF PEER REVIEW EXPERIENCES AND ATTITUDES

To address this gap in the literature on co-reviewing and ghostwriting, we designed a survey to evaluate the frequency of, and rationales for, ghostwriting and co-reviewing by ECRs. The IRB-approved, online survey garnered 498 responses over a month-long data collection period in September, 2018 (<u>Methods: Survey of peer review experiences and attitudes; Appendix: Text of The Role of Early Career Researchers in Peer Review – Survey</u>). Respondents hailed from 214 institutions that were geographically diverse both within and beyond the US. Most participants were from institutions in North America (n = 370), followed by Europe (n = 87) and Asia (n = 21). 74% of all respondents were based in the US, of which 64% were Citizens or Permanent Residents and 36% held temporary visitor status. Institutions from 40 US states or territories were represented, with the most respondents coming from Washington University in St. Louis, University of Kentucky, Rockefeller University, and the University of Chicago (<u>Appendix: List of institutions with multiple survey respondents</u>).

The majority of survey respondents (65%) were ECRs in the life sciences (Figure 1). The five largest groups of self-identified fields were: Neuroscience; Biomedical; Biology; Biochemistry; Cell and/or Developmental Biology (<u>Appendix: List of topics assigned to fields of study</u>). This was as expected given our efforts to primarily engage ECRs and our connections to biomedical postdoctoral populations (<u>Methods: Survey distribution, limitations, and future directions</u>). We surmise that postdocs (63% of all respondents) are over-represented in this survey (although it is difficult to determine the proportions of researchers by career stage in the U.S., particularly as the number of postdoctoral researchers in the US is currently unknown (Pickett et al. 2017)).

**Figure 1:**
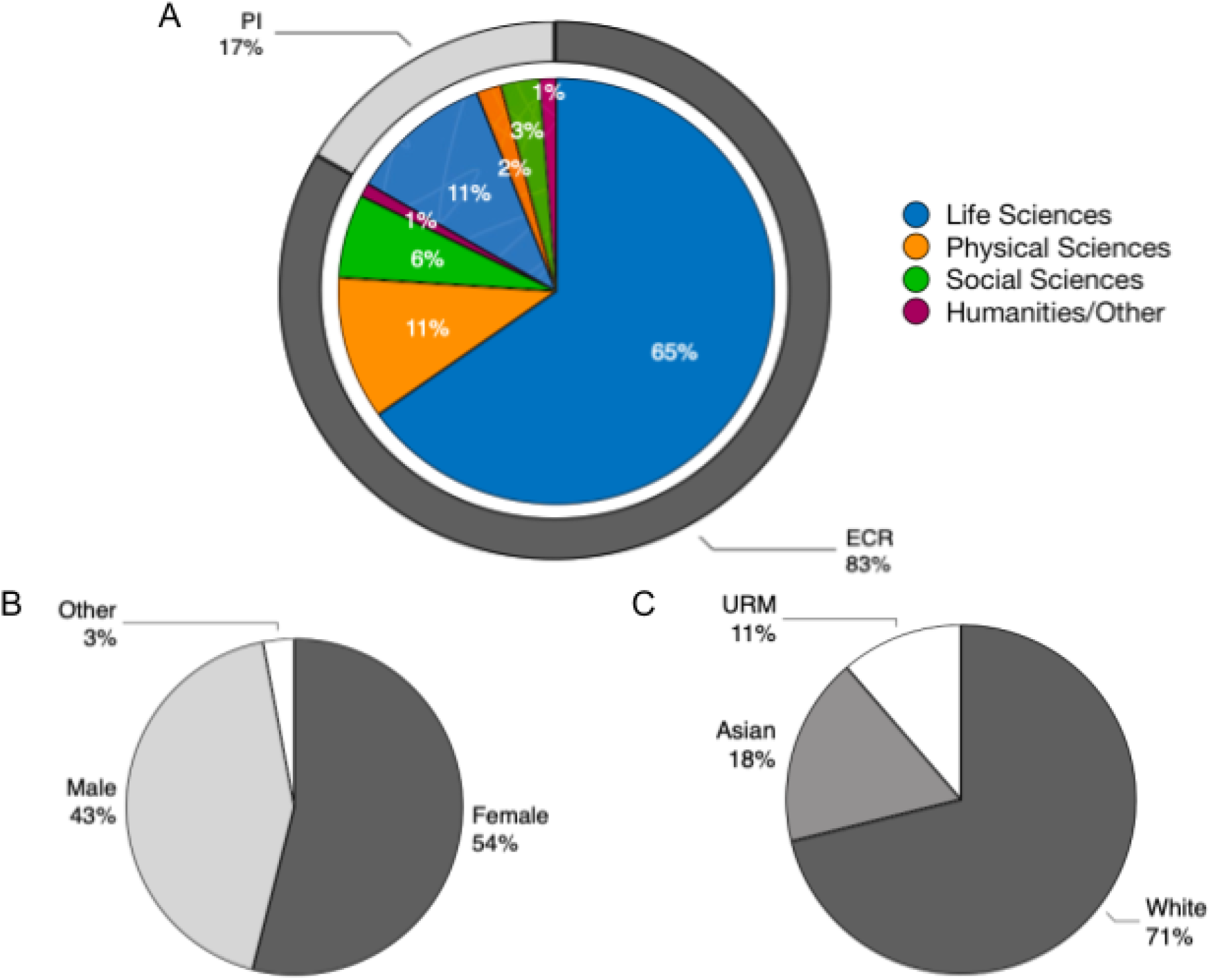
Demographics of survey respondents. (A) Distribution of responses by field of study and career stage. Respondents’ write-in fields of study were categorized for analysis purposes (<u>Appendix: List of topics assigned to fields of study</u>). (B) Distribution of responses by gender. (C) Distribution of responses by race/ethnicity. URM, underrepresented minority in the sciences.

### Ghostwriting happens frequently, despite a common belief that it is unethical

#### Frequency of co-reviewing

The results of our survey support the hypothesis that co-reviewing of manuscripts by ECRs is widespread and frequently goes unacknowledged to journal editorial staff. 73% of *all* survey respondents have acted as co-reviewers and often at numerous times (33% have co-reviewed on 6-20 occasions and *4%* on more than 20 occasions, Figure 2A). Co-reviewing by ECRs is common, with 79% of postdocs and 57% of PhD students having “contributed ideas and/or text to peer review reports where [they are] not the invited reviewer (e.g. the invited reviewer is the PI for whom you work)” (Figure 2B). These data suggest that **collaboration on peer review reports, especially by ECRs who are not the invited reviewer, is an academic norm.** By contrast, when asked about independent reviewing experiences, 37% of *all* survey respondents stated that they had never had such an experience. 35% reported having had this experience on 1-5 occasions; 20% on 6-20 occasions; and 8% on more than 20 occasions. 55% of the ECR respondents have never carried out independent peer review as the invited reviewer (Figure 2C).

**Figure 2:**
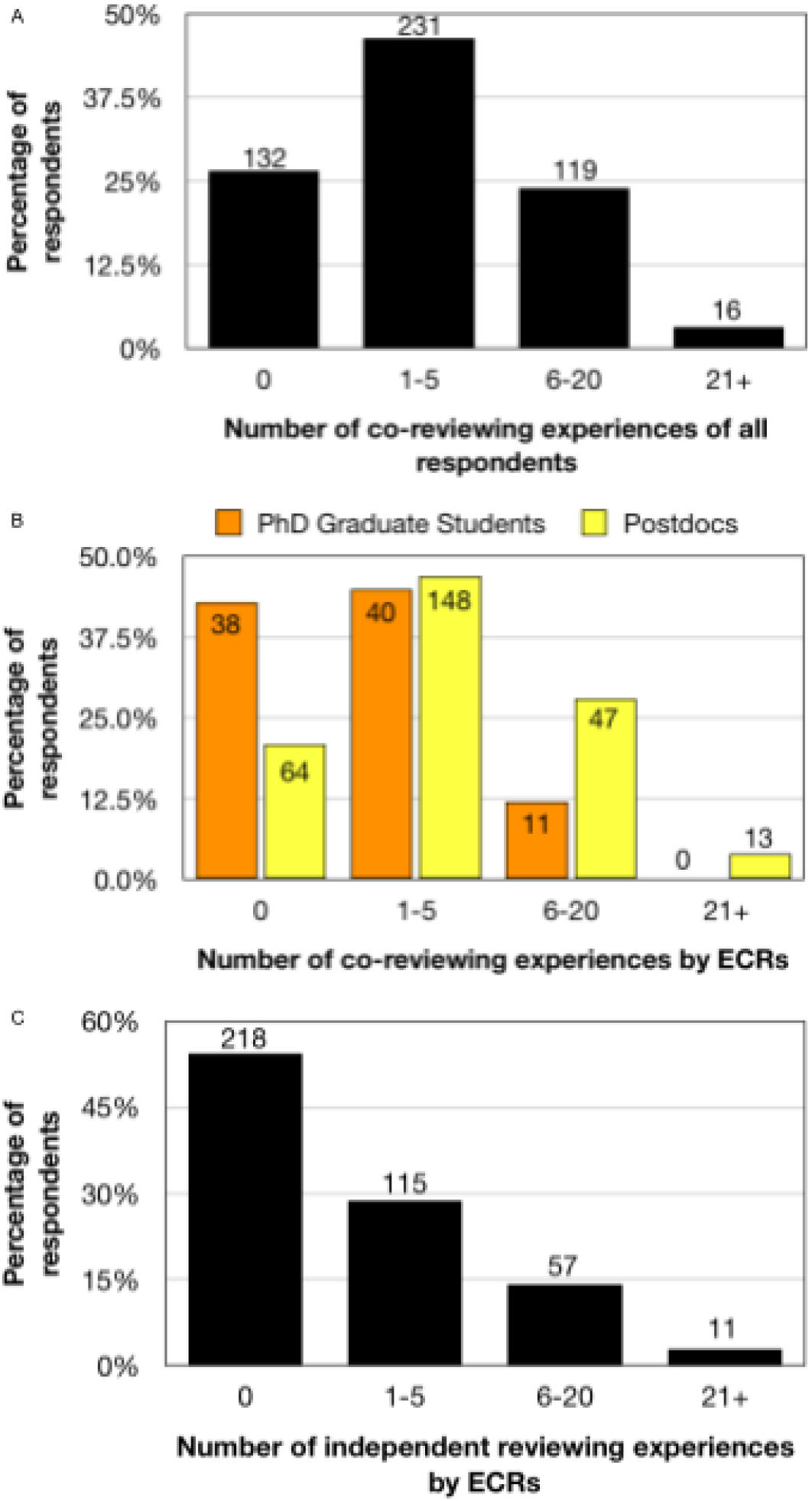
Responses to “How many times in your career have you contributed ideas and/or text to peer review reports where you are not the invited reviewer (e.g. the invited reviewer is the PI for whom you work)?” (A, B) and “How many times in your career have you reviewed an article for publication independently, i.e. carried out the full review and been identified to the editorial staff as the sole reviewer?” (C). (A) Number of co-reviewing experiences of all survey respondents (n = 498). 73% of respondents had participated in co-reviewing, of which 63% had carried out co-reviewing activities on 1-5 occasions; 33% on 6-20 occasions; and 4% on more than 20 occasions. (B) Number of co-reviewing experiences by ECR career stage. The distribution of postdocs (n=312) is skewed toward more co-reviewing experiences while the distribution of PhD students (n=89) is skewed toward fewer co-reviewing experiences. (C) Number of independent review experiences of ECRs. Breaking this down by career stage, we found that 218 respondents (55%) from our pool of 401 ECRs had never carried out independent peer review, and conversely 183 (46%) had carried out independent review as the invited reviewer. Of those who had carried out independent review, 63% (n=115) had done so 1-5 times; 31% (n=57) had done so 6-20 times, and 6% (n=11) had done so more than 20 times.

#### Motivations for co-reviewing

We hypothesized that a significant motivation for ECRs to engage in co-reviewing was to gain experience in peer review of manuscripts, a fundamental scholarly skill. We asked all survey respondents what training they received in peer review of manuscripts (Figure 3). Respondents report that their Pis provide the second most common source of training in peer review, bested only by the passive form of learning “from receiving reviews on my own papers.” Training appears to be a major driver for why ECRs are involved in the peer review of manuscripts and indeed training through co-reviewing was the subject of many publications uncovered by our literature review (<u>Appendix: Literature on ECR involvement in peer review as a training exercise</u>).

**Figure 3:**
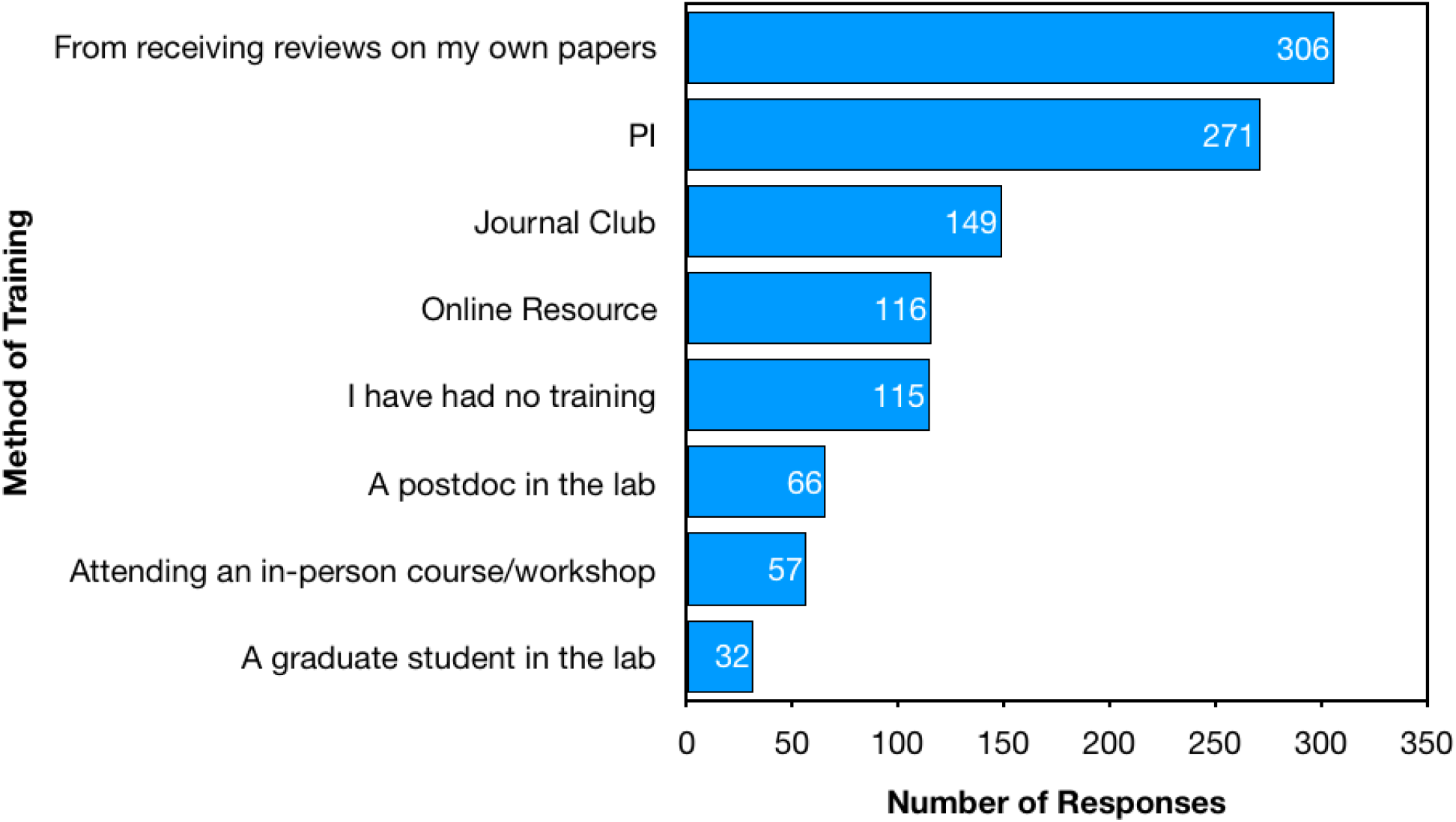
Responses to “How did you gain training in how to peer review a manuscript?” Respondents were able to select as many options as applied to them. These data include responses from all survey participants, including those without any independent or co-reviewing experience.

#### Frequency of ghostwriting

Are these co-reviewers being named to the journal? Or put another way, are journals being made aware that more than just the invited reviewer is contributing ideas and/or text to peer review reports? When we asked “To your knowledge, did your PI ever withhold your name from the editorial staff when you served as the reviewer or coreviewer?,” 46% of respondents knew that their name had been withheld (Figure 4). These data align well with results from a separate question about co-reviewing and ghostwriting experiences: “When you were not the invited reviewer, what was the extent of your involvement in the peer review process?”, to which 44% of respondents reported having had the experience of ghostwriting: “I read the manuscript, wrote the report, my PI edited the report and my PI submitted report with only their name provided to the editorial staff” (Table 1). Taken together, these data suggest that approximately **1 in 2 survey respondents has engaged in ghostwriting of a peer review report on behalf of their PI, the invited reviewer.**

**Figure 4:**
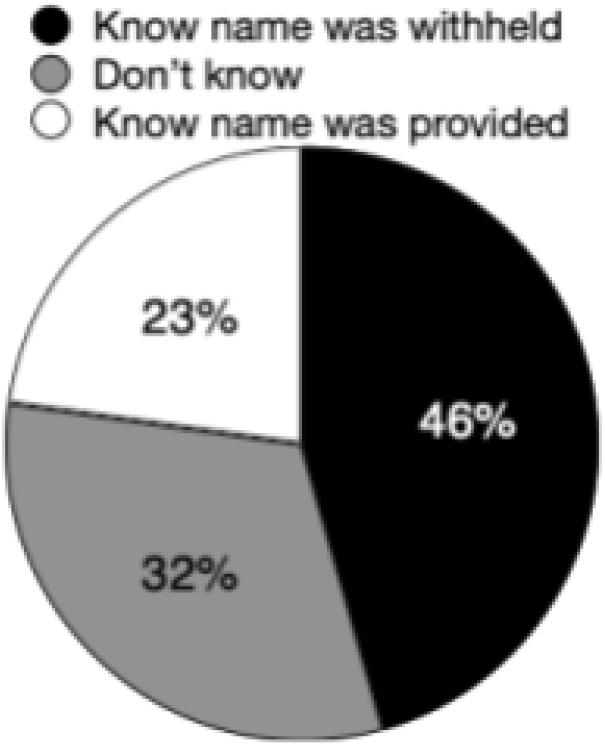
Responses to “To your knowledge, did your PI ever withhold your name from the editorial staff when you served as the reviewer or coreviewer?” 46% of respondents knew that their name had been withheld and 32% did not know. The remaining 23% responded that they knew for certain their name had been disclosed.

**Table 1:** Responses to “When you were not the invited reviewer, what was the extent of your involvement in the peer review process?” Survey participants were able to choose any and all applicable responses from a provided set of possible responses that can be broken down into three interpretation groups. Because respondents were able to select more than one answer, these data include all of the different co-reviewing experiences for each participant.

| Possible survey responses | Respondents that selected this as an answer (%) | Interpretation of response | Respondents that selected at least one of the answers in this group (n, %) |
| --- | --- | --- | --- |
| “I read the manuscript, shared short comments with my PI, and was no longer involved” | **40** | No significant contribution | 149 respondents (**40%**) selected this response |
| “I read the manuscript, wrote a full report for my PI, and was no longer involved” | **47** | Significant contribution, <u>without</u> known credit | 258 respondents (**70%** of those with co-reviewing experience) selected at least one of the responses in this category |
| “I read the manuscript, wrote the report, my PI edited the report and my PI submitted report with only their name provided to the editorial staff” | **44** |  |  |
| “I read the manuscript, wrote the report, my PI edited the report and we submitted the report together with both of our names provided to the editorial staff” | **20** | Significant contribution, <u>with</u> known credit | 80 respondents (**22%** of those with co-reviewing experience) selected at least one of the responses in this category |
| “I read the manuscript, wrote the report, and submitted it independently without my PI’s name provided to the editorial staff” | **3** |  |  |

Note: (Mis)representation of authorship on any scholarly work can be a subjective grey area. We sought to specifically avoid this in our survey questions by using the answers to the question “When you were not the invited reviewer, what was the extent of your involvement in the peer review process?” to disambiguate the grey areas of authorship. We consider any experience that began with “I read the manuscript, wrote a full report for my PI, and...” to be an unequivocally significant contribution deserving of authorship on the peer review report.

Furthermore, of respondents who co-reviewed, 70% report the experience of making significant contributions to a peer review report without knowingly receiving credit (Table 2). These data reveal a breakdown in communication between invited reviewers and co-reviewers. In a more specific follow up question that asked “To your knowledge, did your PI ever submit your reviews without editing your work?”, 52% of our survey respondents report that they were not involved in any editing process with their PIs (Figure 5).

**Figure 5:**
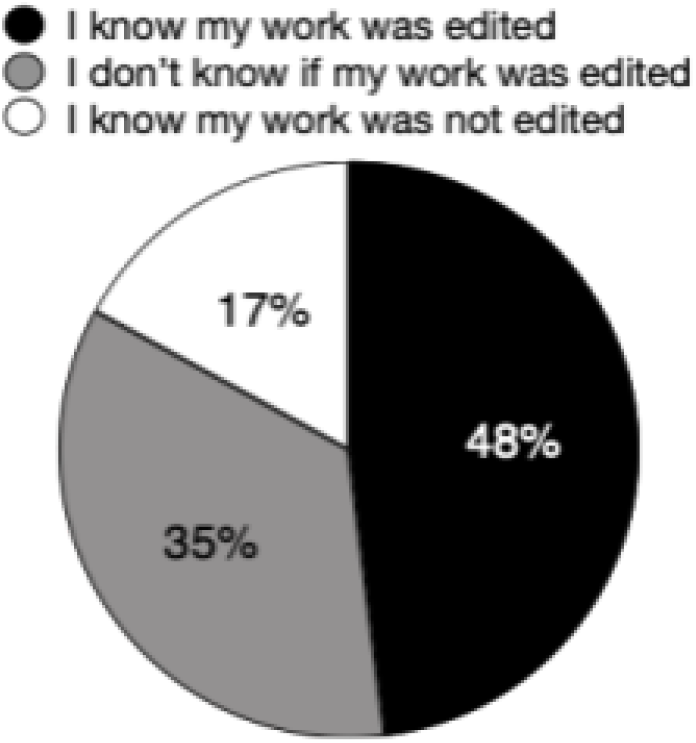
Responses to “To your knowledge, did your PI ever submit your reviews without editing your work?.” More than half of respondents are not involved in editing subsequent to submitting work to the invited reviewer. 17% of respondents answered “yes”, that they knew that their work had not been edited by the PI prior to submission to the journal. Another 35% of respondents were unaware of whether their work was edited by their PI prior to their PI submitting it to the journal. 48% of respondents indicated that “no”, indicating that they knew their work had been edited for sure.

**Table 2:** Statistically significant difference in agreement between ECR and PI populations on Agree/Disagree statements. The higher the mean value calculated for the group, the closer the group feels to disagreeing with statement. “No opinion” responses, coded as 3, are included in these means. A 2-tailed student’s t-test for equality of the means was used to calculate p values. *Indicates that p value was calculated assuming equal variance according to Levene’s test for Equality of Variances.

| **Statement** | **ECR Mean Score** | **PI Mean Score** | **p value** | **% ECRs with no opinion** | **% PIs with no opinion** |
| --- | --- | --- | --- | --- | --- |
| Involving members of a research group in peer review is a beneficial training exercise. | 1.32 ± 0.03<br>(n = 405) | 1.54 ± 0.10<br>(n = 81) | $p = 0.033$ | 2.5 | 2.5 |
| It is ethical for the invited reviewer (e.g. PI) to involve others (e.g. their trainees) in reviewing manuscripts.* | 2.06 ± 0.06<br>(n = 406) | 2.37 ± 0.14<br>(n=81) | $p = 0.029$ | 11 | 15 |
| It would be valuable to have my name added to a peer review report (e.g. to be recognized as a co-reviewer by the editor; or to use a service such as Publons to be assigned credit). | 1.71 ± 0.05<br>(n = 405) | 2.11 ± 0.13<br>(n = 81) | $p = 0.003$ | 10 | 21 |

As mentioned above, an INSIDE *eLife* survey found a slight majority of all respondents had performed peer review with no involvement from their supervisor (Inside eLIFE 2018). The *eLife* survey asked “Have you reviewed before?” and then “If so, to what extent was your supervisor involved?.” Of all respondents in the *eLife* survey (n = 264, 74% of which were postdocs or PhD students), slightly more than half replied “not at all” to the latter question, suggesting that they had done peer review without oversight. One interpretation of these data is that half of respondents had engaged in independent peer review as the invited reviewer. Another interpretation of these data is that half of respondents had carried out peer review with no feedback from their supervisor, the invited reviewer. Our data expand upon that survey by disentangling these two possibilities, asking respondents to reflect on instances “When you were not the invited reviewer.” Now with this added modifier, we still find that **slightly more than half of respondents have written peer review reports without feedback from their PI when the PI is the invited reviewer.**

#### Ethics of ghostwriting and co-reviewing

It seems incongruent and inefficient that half of co-reviewers are not involved in any editing process with their PIs, if a rationale for co-reviewing by ECRs is to receive training in the scholarly activity of peer review. At best, writing a peer review report without receiving feedback from one’s PI prior to submission to a journal is a missed opportunity for the co-reviewer to receive training in this critical skill. At worst, a peer review report that is written by one person and submitted to the journal under the name of another person (the PI) might be considered a misrepresentation of authorship and so a breach of academic integrity. The Office of Research Integrity (ORI), which oversees Public Health Service-funded research in the biomedical sciences, defines plagiarism according to the American Association of University Professors wording as “taking over the ideas, methods, or written words of another, without acknowledgment and with the intention that they be taken as the work of the deceiver” (American Association of University Professors 1989). More specifically, the ORI states that “academic or professional ghost authorship in the sciences is ethically unacceptable”… “because the reader [in this case, the journal editorial staff] is misled as to the actual contributions made by the named author” (Citation: Guideline 27; https://ori.hhs.gov/plagiarism-34).

##### Our survey respondents strongly agree with the ORI, and specifically believe that ghostwriting of peer review reports is an unethical practice

83% of respondents disagree with the statement that “The only person who should be named on a peer review report is the invited reviewer, regardless of who carried out the review”; 81% disagree with the statement that “Ghostwriting a peer-review report for your PI is an ethically sound scientific practice” and 77% disagree that “It is ethical for the invited reviewer (e.g. PI) to submit a peer review report to an editor without providing the names of all individuals who have contributed ideas and/or text to the report” (Figure 6). Put another way, 74% agree that “Anyone that contributes ideas and/or text to the review report should be included as a co-author on the review.” Sharing co-reviewer names with the journal staff is considered not only ethical, but also valuable, with 82% agreeing that “It would be valuable to have my name added to a peer review report (e.g. to be recognized as a co-reviewer by the editor; or to use a service such as Publons to be assigned credit).” Similarly, co-reviewing itself is considered ethical and valuable, with 73% agreeing that “It is ethical for the invited reviewer (e.g. PI) to involve others (e.g. their trainees) in reviewing manuscripts” and 95% agreeing that “Involving members of a research group in peer review is a beneficial training exercise.” The latter statement evoked the strongest positive sentiment of all 11 possible Agree/Disagree statements.

**Figure 6:**
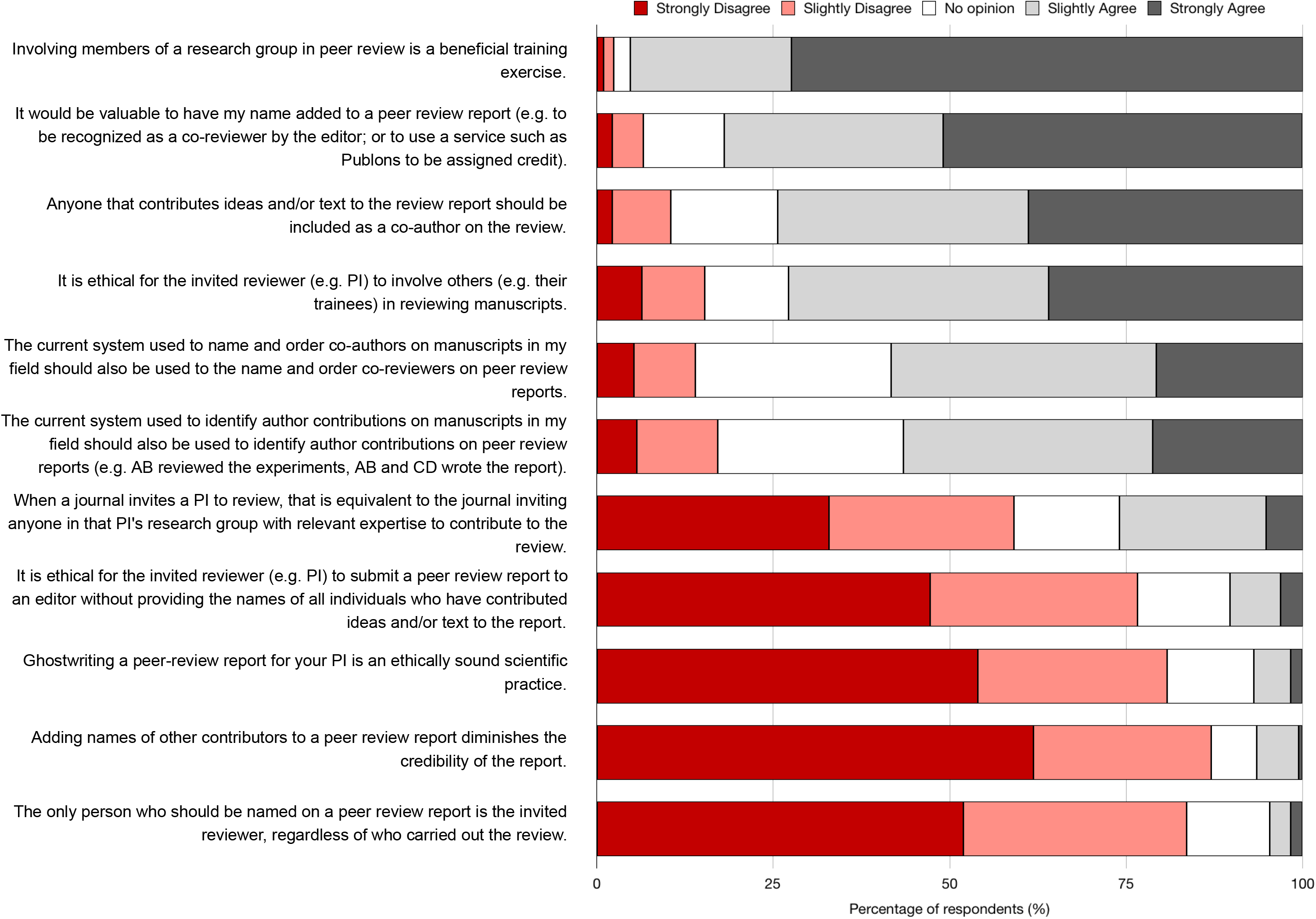
Responses to “Please indicate how strongly you agree with the following statements.” Data represent the opinions (not experiences) of all respondents regardless of whether or not they had participated in peer review. Respondents were also provided with a textbox to submit comments to expand and/or clarify their opinions.

Since ECRs are the disadvantaged population when ghostwriting occurs, we hypothesized that ECRs might have more strongly held opinions about these practices than PIs. We calculated the degree to which ECRs and PIs agreed with each other by setting 1 as Strongly Agree through to 5 as Strongly Disagree, and 3 set as No Opinion, and then by calculating the mean degree of agreement as a score. Overall, there was a great degree of similarity between the responses of ECRs and PIs. We found only 3 of 11 statements where there was a significant difference in the extent of agreement between ECRs and PIs (Table 2). In all of these cases, ECRs felt significantly more strongly than PIs but still shared the same valance (e.g. both groups agreed or both groups disagreed with the statement, just to a differing strength). Since 21% of PIs vs. 10% of ECRs responded with “no opinion” to the statement about the value of credit for co-reviewing, we wondered whether the difference in means for this statement could be ascribed to more indifference on the part of the PI population. We recalculated the means removing the “no opinion” responses and found that the difference between the means was still significant but much less so (p = 0.048; ECRs: 1.57 ± 0.05 (n = 365) vs. PIs: 1.88 ± 0.15 (n = 64)).

#### Motivations for ghostwriting

##### If 4 out of 5 survey respondents think ghostwriting is unethical, then why do half of all respondents participate?

We measured the motivations for ghostwriting by: (1) asking all survey respondents, regardless of peer review experience, to surmise why someone might withhold the name of a co-reviewer (Figure 7), and (2) asking only survey respondents with ghostwriting experience to report specific reasons that the invited reviewer gave for withholding their name (Table 2). In this way, we compared cultural beliefs with actual practice.

**Figure 7:**
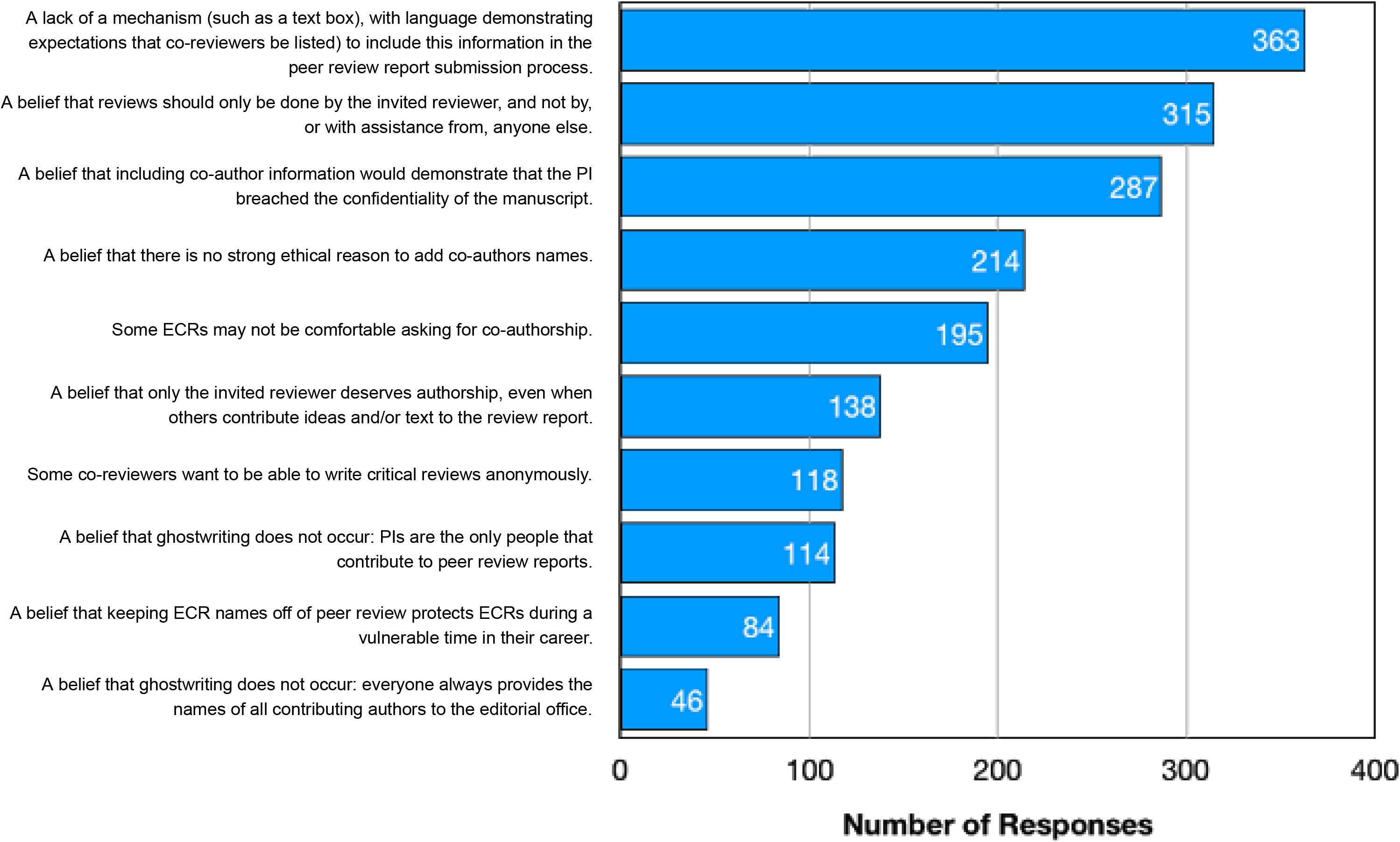
Responses to “What do you think are the reasons why the names of co-authors on peer review reports may not be provided to the editorial staff?.” Here our intent was to ask the respondents *not for their personal opinions on whether co-reviewers should be named*, but rather for their perspective on what they thought the logistical or cultural barriers may be that would cause names to be withheld in general practice. Respondents were able to select as many answers as they felt applied. The frequencies do not allow us to assess how important the barriers are, and respondents were not asked to rank barriers, but simply to surmise which ones they felt were relevant to the current practice of ghostwriting.

The most commonly-believed barrier to naming co-reviewers was a lack of a physical mechanism to supply the name to the journal, with 73% of respondents selecting this as an option (Figure 7). Cultural expectations were the next most commonly-cited barriers, such as “A belief that reviews should only be done by the invited reviewer, and not by, or with assistance from, anyone else” and “A belief that including co-author information would demonstrate that the PI breached the confidentiality of the manuscript”, selected by 63% and 58% of respondents respectively. These latter responses allude to journal policies prohibiting invited reviewers from sharing unpublished manuscripts without prior permission. All three top responses relate to journal policies that are either absent or prohibitive. Write-in responses, summarized in Box 2, echo themes about how ghostwriting is simply the status quo in peer review. At the same time, respondents also wondered why this is the case, and why including co-reviewer names is not common practice.

###### BOX 2: THEMES FROM WRITEIN REASONS FOR WHY GHOSTWRITING MAY OCCUR

**Box 2:** Themes and supporting examples of write-in responses to the question: “What do you think are the reasons why the names of co-authors on peer review reports may not be provided to the editorial staff?”

| BOX 2: THEMES FROM WRITE-IN REASONS FOR WHY GHOSTWRITING MAY OCCUR |  |
| --- | --- |
| Theme | Example write-in responses |
| Cultural expectations | "A belief that ghostwriting does occur, but everyone accepts that it's just the way it is." |
|  | "The belief it has always been like this so why doubt /change the process" |
|  | "PIs simply don't think of it because they're used to doing things this way" |
|  | "PIs think this practice is okay." |
| Training | "A belief that this is 'how it is done,' and inviting trainees to contribute to reviews is important for their training, but it is not necessarily important for them to get credit for it." |
|  | "PI feels while the ECR is being trained in doing the review should not be listed as co-author of the review." |
| "I don't understand it" | "Either as a reviewer or as an editor, I would have no problem with a co-review. I'm not really sure why more people don't do it. They absolutely should." |
|  | "I have no idea why this is not common practice" |

In contrast, when we asked “Consider cases where you contributed to a peer review report and you know your name was NOT provided to the editorial staff. When discussing this with your PI, what reason did they give to exclude you as a co-reviewer?” 73% of respondents reported that they had not discussed this with their PI (Table 3). This is consistent with the lack of communication between invited reviewers and co-reviewers documented above (Table 1, Figure 5). Of the 27% of respondents who had ghostwritten and *did discuss* the matter with their PI, most were told that the reason their name was withheld was either a prohibitive journal policy and/or prevailing cultural expectations about co-authorship on peer review reports. Only *4%* of those who had discussed the matter with their PI cited a practical barrier, such as the lack of a text box for co-reviewer names on the journal review submission form, as the reason given by their PI for withholding their name. Write-in responses to this question again refer to cultural expectations as the major drivers for ghostwriting (Box 3; <u>Appendix: Written responses giving reasons PIs excluded survey respondents as co-reviewers</u>). Many of these write-in reasons articulate that it is good practice for ECRs to participate in peer review; however, they simultaneously fail to explain why this necessitates withholding the names of co-reviewers. These data suggest that there is a common conflation of ghostwriting (withholding names) with co-reviewing (involving ECRs in peer review, often for the purposes of training).

**Table 3:**
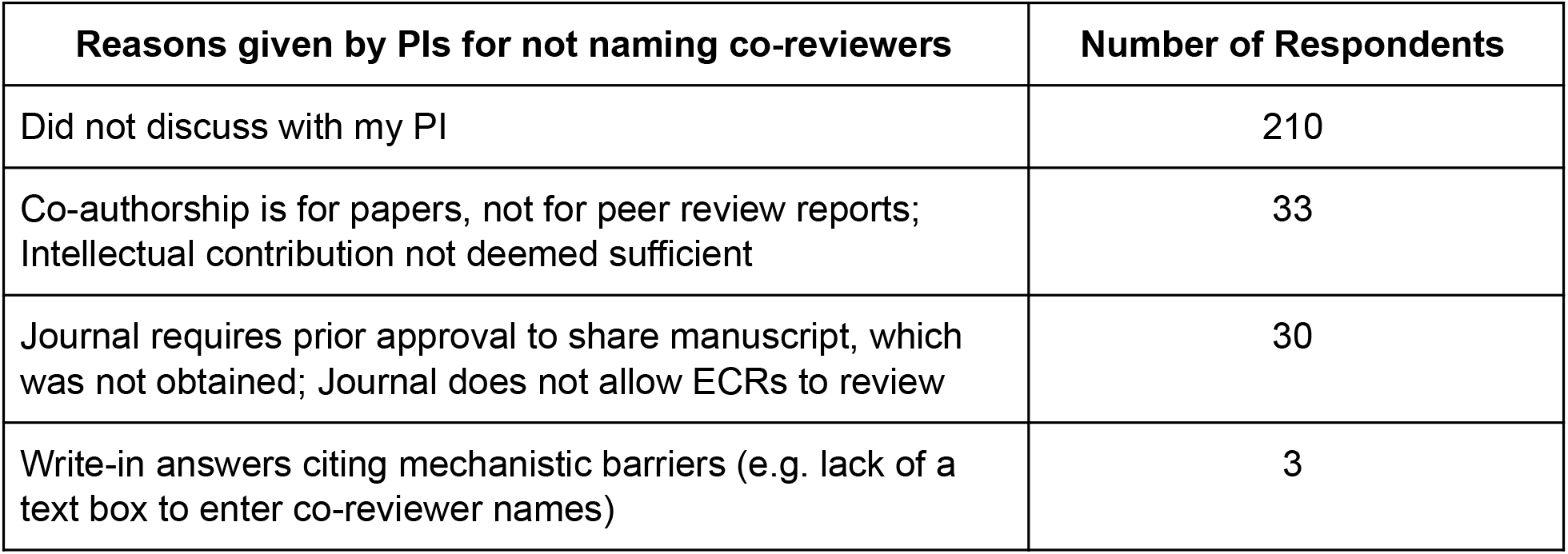
Responses to “Consider cases where you contributed to a peer review report and you know your name was NOT provided to the editorial staff. When discussing this with your PI, what reason did they give to exclude you as a co-reviewer?” In addition to the possible answers provided by the survey, respondents were also provided with a textbox to add write-in responses.

| Reasons given by PIs for not naming co-reviewers | Number of Respondents |
| --- | --- |
| Did not discuss with my PI | 210 |
| Co-authorship is for papers, not for peer review reports; Intellectual contribution not deemed sufficient | 33 |
| Journal requires prior approval to share manuscript, which was not obtained; Journal does not allow ECRs to review | 30 |
| Write-in answers citing mechanistic barriers (e.g. lack of a text box to enter co-reviewer names) | 3 |

###### BOX 3: THEMES FROM WRITEIN REASONS GIVEN BY PI FOR WITHHOLDING ECR NAME

Themes and supporting examples of write-in responses to the question “Consider cases where you contributed to a peer review report and you know your name was NOT provided to the editorial staff. When discussing this with your PI, what reason did they give to exclude you as a co-reviewer?”

| BOX 3: THEMES FROM WRITE-IN REASONS GIVEN BY PI FOR WITHHOLDING ECR NAME |  |
| --- | --- |
| Theme | Example write-in response |
| Sin of omission | “They forgot” |
|  | “Didn’t think of including me; didn’t know how to do so” |
|  | “He was in a hurry and he couldn’t figure out the journal’s website” |
| Cultural expectations | “This was not explicitly discussed, but the PI implied this is “common practice” and normal for ECRs to gain experience” |
|  | “[PI] said only [they] would be invited to review for such a prestigious journal and “we” need this for future submissions” |
|  | “Apparently this duty is part of my job description” |
|  | “I was told this is how one gets to train to review papers and grants” |
| A good way to train | “Reviewing papers as [an] ECR is part of the ECR training” |
|  | “It’s good experience for me.” |
|  | “It was good for my career to practice” |

## DISCUSSION

Journal peer review is an important part of scholarly work and, as such, scholars should be well-trained in this activity to ensure quality peer review into the future. Our dataset demonstrates that involving ECRs in peer review is an academic norm, with ¾ of our survey population having contributed significantly to a peer review report when they were not the invited reviewer (co-reviewed) and ½ having done so without being named to the journal editorial staff (ghostwritten). These high frequencies starkly contrast with journal policies and widely-held cultural expectations that only the invited reviewer engages in the peer review of a manuscript for publication. They also fly in the face of our values as a community, when 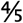 of those surveyed agree that ghostwriting is unethical. What drives these differences between community values and experience, and what can be done to reconcile them?

### Explanations for ghostwriting are conflated with explanations for co-reviewing

Co-reviewing as a training exercise and ghostwriting are separable processes: training through co-review can and does happen *whether or not* named credit is given to the co-reviewer, and excluding co-reviewer names from peer review reports can and does happen *whether or not* the co-reviewer has experienced quality training in the process. Even as we sought to collect data that would disentangle these two processes, we found that the rationales for ghostwriting were all too often conflated with the rationales for co-reviewing. For example, when we asked respondents specifically for the reason(s) their PI gave them for excluding their names on a peer review report they co-authored, many wrote-in responses such as “I was told this is how one gets to train to review papers…” (Box 3). This explanation for why ghostwriting occurs – that it is a beneficial and common practice for ECRs to participate in peer review as a training exercise – does not actually answer why co-reviewers are not named to the journal editor. **An important first step in reducing unethical ghostwriting practices will be to decouple it in the Zeitgeist from the beneficial training of ECRs through the process of co-review.** We therefore separate our discussion of the motivations for co-reviewing and ghostwriting in an effort to find solutions to ghostwriting that simultaneously support ECR co-reviewing as training in peer review.

### Whv does co-reviewina happen so frequently?

#### Co-reviewing by ECRs as a common and valuable training exercise

Perhaps the most intuitive and optimistic answer to the question stated in the heading is that, when the issue of named credit on peer review reports is set aside, **co-reviewing by ECRs is considered a valuable, ethical, and common form of training in the peer review of manuscripts.** 95% of survey respondents find co-reviewing to be “a beneficial training exercise” and 73% agree that “it is ethical for the invited reviewer (e.g. PI) to involve others (e.g. their trainees) in reviewing manuscripts.” Co-reviewing is also the second most commonly reported source of training in peer review, bested only by “receiving reviews on my own papers,” a method of passive learning. These data should be weighed heavily when considering journal policies regarding co-reviewing broadly, and ECRs as co-reviewers specifically, since **any policy that prevents co-reviewing by ECRs would remove a common and valuable training exercise in peer review.**

Survey respondents clearly find that co-reviewing by ECRs has significant benefits and is not inherently problematic. Yet problems with using “ECR training” as a rationale for co-reviewing arise when training doesn’t actually take place and instead co-reviewing devolves into an exploitative delegation of scholarly labor, when this mode of training is ineffective at improving ECR’s ability to peer review, and/or when co-reviewing perpetuates unethical conditions like ghostwriting or breaches of manuscript confidentiality.

#### Co-reviewing by ECRs as a delegation of scholarly labor

A major limitation with using “ECR training” as a rationale for co-reviewing arises when half of survey respondents report writing a peer review report without any training interaction with their PIs. The hidden delegation of scholarly labor to ECRs benefits the invited reviewer, often an overburdened PI in a hypercompetitive research environment (Alberts et al. 2014). As the number of ECRs has grown steadily in biomedicine since the mid-20th Century, academic positions to employ them have not (Heggeness et al. 2016; Heggeness et al. 2017). The resulting hypercompetitive environment incentivizes the use of ECRs as cheap labor to fulfil productivity requirements. This can include scholarly labor that PIs lack the time or even expertise to perform, such as manuscript peer review.

These market forces provide a second explanation, beyond ECR training, for why co-reviewing is commonplace: that the selective pressures of a hypercompetitive research environment have caused co-reviewing and ghostwriting by ECRs to become the status quo. Our survey respondents agree, with many providing write-in responses such as: “apparently this duty is part of my job description” (Box 3). The delegation of scholarly labor to ECRs is not necessarily, nor intentionally, exploitative, although it can easily become so given the power dynamics and documented lack of communication between mentors and mentees (Van Noorden 2018). Any successful intervention to address concerns about the ethics of co-reviewing by ECRs should take into account this status quo – that **it is commonly an unstated expectation that ECRs carry out peer review on behalf of their PI, and that ECRs may not feel they have the freedom to decline even if they feel it is unethical to participate.** We encourage the scientific community to recognize this dynamic and recommend that journals adopt policies that specifically address this common yet unethical practice.

### Limitations to depending on co-reviewinα as training

#### A lack of evidence-based training structures and community oversight

The top three forms of peer review training reported by our survey are notable in that they do not necessarily involve any formal training structure, quality control, or community oversight (Figure 3). “Receiving reviews on my own papers” is only able to give a limited number of examples of how others review. Since it is a common complaint that reviews are overly critical and fail to give constructive feedback (Schneiderhan 2013), it seems counter-intuitive to have this be the main example by which ECRs learn how to conduct peer review. This is also a passive form of learning that lacks individualized or iterative feedback from a mentor. By contrast, training received “from PI” may benefit from a personalized teaching relationship; however, the quality of this teaching is again dependent on the training the PI may have received, which may be from their own reviews and previous mentors. Journal clubs too may be self-organized groups comprised of e.g. only graduate students, that may lack a desired level of peer review expertise and/or a rigorous training component.

These data suggest that **one’s training in peer review is largely determined by a small number of individual experiences outside of evidence-based training structures.** Such training experiences are likely to be self-reinforcing, their quality dependent on the quality of the reviews received or the quality of training received by PIs themselves. The small sample size of these experiences may also reinforce a number of biases, including selection bias where one’s few experiences may not be representative of the population, memory biases where negative experiences (e.g. from receiving highly critical reviews) are more readily remembered and so taught (Kensinger 2007), and gender or other demographic biases currently being studied in the peer review process (Murray et al. 2018). The quality and consistency of training that is drawn from a small number of different personal experiences, in turn, is likely to be highly variable.

These results highlight an area of opportunity to improve and standardize training in peer review of manuscripts as a critical scholarly skill. One intervention to be considered is the establishment of more standardized, evidenced-based training for peer review of manuscripts. This might be accomplished by graduate programs, scientific societies, or even journals themselves, and some already provide training materials (<u>Appendix: Examples of formalized training in peer review</u>). Given the central place that peer review has in maintaining trust in our ability to scrutinize each other’s work (Patel et al. 2017) and the public’s trust in our enterprise, we are concerned that insufficient attention is given to rigorous peer review training from the very beginning of graduate programs. That “Peer Review 101” does not appear to be a compulsory and ubiquitous course at graduate schools should give the community pause for thought, and future efforts should tie mentoring efforts with evidence-based training in the peer review process to ensure that all PhD-holders are competent and capable of engaging in a constructive peer review process.

#### A lack of training the trainers

Since the top two reported forms of training involve PIs (either as manuscript reviewers or ECR mentors), interventions that attempt to ensure PIs have received training in peer review *and in communicating this skill to ECRs* may also be appropriate. We have encountered the perspective that a postdoc should not undertake peer review while a PI is assumed to be expert at it, and **this is illogical, given that there are no systematic steps to ensure proficiency at peer review during the postdoc-to-PI transition.** Indeed, a lack of “training the trainers” was cited as a main reason for why pairing experts with new peer reviewers failed to improve review quality in one of the few randomized controlled trials of this practice (Houry et al. 2012). In this study, the expert reviewers were not given any training themselves in an effort to minimize their burden, which the authors conclude resulted in an ineffective training program. They suggest that effective mentoring of new reviewers should include training in providing feedback from expert to trainee. Others echo this recommendation, finding that iterative feedback from the mentor leads to positive outcomes as measured by reviewers’ beliefs and by objective evaluations of improved review quality (Castelló et al. 2017; Doran et al. 2014). One mechanism that proved efficacious was directing the PI to report on the same manuscripts as the trainees, thereby providing trainees with an example/comparison peer review report (Castelló et al. 2017). An emerging theme, therefore, is be **a need for improved communication between mentors and mentees during the co-review training experience.**

### Whv does ghostwriting happen so frequently?

#### A lack of communication about authorship of peer review reports

Even in the best case scenario for co-reviewing as a training exercise, when training *is* taking place and *is* effective, its benefits can still be confounded by ethical lapses such as ghostwriting. The most common explanation for ghostwriting was that providing co-reviewer names to the editor was simply not discussed (Table 3, 73% of responses from those who knew their names had been withheld). Of those who had contributed significantly to peer review reports with or without known credit, 39% of their experiences fall into the category of “I read the manuscript, wrote a full report for my PI, and was no longer involved” (n = 107 of 274 total responses that begin with “*I read the manuscript, wrote the full report…*”, Table 1). These **data reveal a significant breakdown in communication between invited reviewers and those actually writing the peer review report,** that is concerning on many levels – e.g. with regards to plagiarism, mentor-mentee relationships, and the exploitation of ECR labor.

Why is authorship on peer review reports not a topic of conversation between collaborating invited reviewers and co-reviewers, as it might be between senior and junior authors on a manuscript? One simple reason may be that not naming co-reviewers is the expected status quo. This theme is echoed even when discussions between PIs and ECRs do occur: 25% of respondents who discussed the issue with their PI were told that the reason their name was excluded was “co-authorship is for papers not for peer review reports” (Table 3). Many write-in responses support this idea that ghostwriting may come about as a “sin of omission:” [PI] “didn’t think of including me” and “they forgot” (Box 3). ECRs express similar opinions, with one postdoc writing: “I’d never really thought about this before. I just assumed it was part of the process. But it is very time consuming and I do believe that all reviewers should receive credit for the review.” Therefore, if ghostwriting arises from PIs and ECRs simply not thinking to include co-reviewer names on peer review reports vs. intentional withholding of names, then building awareness about this issue should encourage more conversation within the community and between mentors and their trainees. Mentors should discuss with mentees how they approach peer review and how credit is given for peer review work to impart proper review ethics to trainees.

We recognize that a naive “sin of omission” may not be the only roadblock that prevents open conversation about authorship between PIs and ECRs. Certainly, power imbalances might prevent an ECR from feeling able or willing to initiate this conversation with their PI. Given that the prevailing cultural norm is to not provide co-reviewer names, ECRs may not feel equipped with evidence to support their self-advocacy, regardless of how strongly they feel about the topic. Indeed, 39% of respondents think that ghostwriting occurs because “some ECRs may not be comfortable asking for co-authorship” (Figure 7). As one write-in states: “you don’t want to piss off the boss.” Ghostwriting, therefore, may be a symptom of the larger problem in academia that ECRs are extremely dependent on their PI’s goodwill for retention in the hypercompetitive research environment (eg. for letters of recommendation throughout their career, or immigration status).

#### A lack of perceived value in naming co-reviewers?

Do co-reviewers see a benefit to having their name disclosed to the journal editor? 82% of survey respondents agree that “It would be valuable to have my name added to a peer review report.” Co-reviewers, especially ECRs, may value having their name provided to the journal for many reasons, including the ability to be “known” in the field to scientific editors and potential colleagues; the ability to have their scholarly work acknowledged by a verified third party service (e.g. Publons) for career advancement; and/or the ability to demonstrate eligibility for visas and residency. While not all co-reviewers will find equal benefit in receiving credit for every co-reviewing experience, **each individual should have the ability to choose to be named as a co-reviewer.** A lack of universal benefit in being named as a co-reviewer is not a suitable reason to withhold co-reviewer names.

It is the co-reviewer and not the invited reviewer who perceives most benefit from having recognized co-authorship of a peer review report. We see in our data that, while PIs and ECRs both agree that there is value in this practice, ECRs agree more strongly than PIs that there is value in receiving credit or being known to the journal editorial staff as a co-reviewer (Table 2). This was one of only 3 (of 11) statements in which there was a significant difference between the mean agreement score of ECRs and PIs. More PIs also had no opinion on this statement than ECRs (21% of PIs vs. 10 *%* of ECRs having no opinion). This may suggest that the difference in strength of agreement is not due to opposition, per se, but rather due to greater indifference on the part of the PI population, who may not see value in it, perhaps because they are already known to editors. In the words of one write-in response: “PI surprised I would be interested in being acknowledged, and seemed like too much trouble to acknowledge my contribution.” This ambivalence for due credit for co-reviewers likely derives from a position of social privilege. Ironically, it is the individuals in the scientific community with positions of relative privilege who may be less likely to recognize the benefit of credit for co-reviewing – e.g. tenured PIs, US citizens – and yet who may have the greatest power to change the status quo. When people of privilege are the only community members that participate in decision-making, they may (inadvertently) create policies or reinforce cultural standards that fail to consider the differing values of less privileged members of the community members, like ECRs. We support the growing effort to include more diverse voices in leadership roles in science, including Future of Research’s “Who’s on board” initiative to seat more ECRs on the advisory boards of scientific organizations and journals, citing examples such as the Early Career Advisory Group at *eLife* (https://elifesciences.org/about/early-career).

#### A perceived value in withholding co-reviewer names?

Survey respondents clearly see a benefit to having co-reviewer names disclosed to the journal editor. At the same time, do survey respondents think that there is any counterbalancing value *to not naming* co-reviewers to the editor? We wondered whether ghostwriting might occur because of the perception that adding co-reviewer names diminishes the peer review report itself (e.g. by providing evidence that someone other than the invited reviewer contributed to the review)? Survey respondents overwhelmingly do not find this to be the case, with 87% of respondents regardless of career stage disagreeing with the statement that “Adding names of other contributors to a peer review report diminishes the credibility of the report.” This statement evoked the strongest negative sentiment of all 11 possible Agree/Disagree statements. We would be curious to know the opinion of journal editorial boards on this subject, since there are journals with policies that prevent ECRs from serving as co-reviewers (or invited reviewers). **If journal policies derive from a concern that ECR contributions to peer review diminish the credibility of the report, then they would be significantly out of line with the scientific community’s opinion represented in this dataset.** They also go against the results of an ongoing experiment in co-reviewing at 3 Elsevier journals, in which editors did not rate the co-reviewed reports as poor or low quality, and more than half were considering co-reviewers to serve as invited reviewers on future manuscripts (Mehmani 2019). Indeed, that there should be no diminution of quality in co-reviewed peer review reports supports research suggesting that reviewers who are earlier in their careers may be perceived by editors as better reviewers (Black et al. 1998; Evans et al. 1993; Callaham and Tercier 2007). Being closer to bench research, rather than having more experience in reviewing itself, may be a key determinant of this trait (Stossel 1985).

Another reason for why there might be perceived value in withholding co-reviewers names to the editor is to protect ECRs during a vulnerable time in their careers. Occasional write-in responses support this hypothesis, such as “It [being named on a peer review report] may give certain ECRs a bad reputation if they review things really harshly.” However, on the whole, survey respondents largely do not see this protectionism argument as a driver of ghostwriting, with only 17% of respondents choosing “a belief that keeping ECR names off of peer review reports protects ECRs during a vulnerable time in their career” as a potential reason for why ghostwriting occurs. This finding might be because we specially asked about traditional peer review, where reviewer names are known only to the journal editor and excluded cases of open peer review, where the names of reviewer(s) might be made more widely available. Moreover, the choice of a contributing author to withhold their name on occasion – as would be the case if a collaborator chose to recuse their name from a manuscript – does not justify the use of protectionism as an explanation for ghostwriting in general. In summary, **survey respondents find added value in naming co-reviewers, and also see no loss of value to peer review reports when co-reviewer names are added.**

#### A lack of a practical mechanism for reporting co-reviewers?

Could it be that a main driver of ghostwriting boils down to a practical roadblock such as a lack of clear reporting mechanism? An obvious, unavoidable mechanism (e.g. a mandatory checkbox and/or textbox) for reporting co-reviewer names on the submission page for peer review reports might be necessary and sufficient to prompt an invited reviewer to discuss co-reviewer authorship and/or to name co-reviewers to the journal editors. We hypothesized that a lack of such a mechanism was a major reason why co-reviewer names are often excluded. Indeed, respondents seemed to agree with our hypothesis, with 73% surmising that ghostwriting occurs because of “A lack of a mechanism (such as a text box, with language demonstrating expectations that co-reviewers be listed) to include this information in the peer review report submission process.”

However, we noticed a striking difference between these beliefs and the actual experiences reported in our survey. When we asked unnamed co-reviewers to reflect on their own experiences with **ghostwriting, only 3 responses cited a lack of a practical mechanism such as a checkbox.** This represents only *4*% of those who were given a reason by their PI for withholding named credit (n = 77). These data reveal a major difference between a culturally perceived barrier and an actual barrier to naming co-reviewers. A policy shift that fixes this practical mechanism – e.g. adding a checkbox – may still help and may feel like the right thing to do but may not be as impactful as hoped if it fails to address the actual barriers that are faced.

Our data reveal two specific barriers to naming co-reviewers on peer review reports, in addition to the more general problem of a lack of communication about authorship. Both of these barriers arise because **common practice is out of alignment with consensus opinion about best practices,** according to survey respondents.

#### A belief that co-reviewers don’t deserve credit regardless of how much they contribute

The first of these rationales for ghostwriting is a cultural expectation that co-reviewers don’t deserve named credit to the editor regardless of how much they contribute. 43% of ghostwriting experiences were explained by “co-authorship is for papers, not for peer review reports” or “intellectual contribution not deemed sufficient” even though the question stipulated to only consider cases where the co-reviewer contributed ideas and/or text to a peer review report. These experiences are substantiated by community perceptions, with 28% of respondents surmising that ghostwriting occurs due to “A belief that only the invited reviewer deserves authorship, even when others contribute ideas and/or text to the review report.” These given rationales for ghostwriting contradict community opinion about whether this *should be* the case, with 83% disagreeing that “the only person who should be named on a peer review report is the invited reviewer, regardless of who carried out the review” and 74% agreeing that “anyone that contributes ideas and/or text to the review report should be included as a co-author on the review.”

#### Prohibitive journal policy that is out of alignment with current practice

The other most commonly reported reason given by invited reviewers to justify ghostwriting experiences to co-reviewers is prohibitive journal policies. 39% of co-reviewers who discussed with the invited reviewer the possibility of including their name on the peer review report cite “journal requires prior approval to share manuscript, which was not obtained” or “journal does not allow ECRs to review” as the reason why PIs chose to withhold their names. These experiences are substantiated by community perceptions, with 58% of respondents surmising that ghostwriting occurs because of “a belief that including co-author information would demonstrate that the PI breached the confidentiality of the manuscript” and 63% surmising that ghostwriting occurs because of “a belief that reviews should only be done by the invited reviewer, and not by, or with assistance from, anyone else.” These are the second and third most commonly selected responses out of 11 provided responses. We interpret this to mean that the community understands that many journals have policies that prevent invited reviewers from sharing manuscripts with anyone else and/or policies that prevent ECRs from serving as reviewers or co-reviewers without prior permission from the editor^2^. Yet it seems that, in practice, invited reviewers may not seek prior permissions or may not wish to risk doing so and so choose instead to not reveal the presence of co-reviewers upon submission. In these cases, adding co-reviewer names to a peer review report is equivalent to admitting that journal policies were disobeyed. Given how frequently ghostwriting occurs based on survey data, and how commonly journal policies are cited as the reason for ghostwriting, it seems that **current policies that require invited reviewers to gain permission of the editor prior to involving ECRs in peer review are not effective deterrents for ghostwriting.** Instead, these policies may have the opposite, if unintended, consequence of preventing invited reviewers from feeling free to disclose co-reviewer names upon submission and so promoting unethical ghostwriting practices.

### A call for harm reduction interventions

Our data suggest that **one unethical practice – ghostwriting – is being justified by another unethical practice that is a pervasive status quo – sharing a manuscript without editorial permission with a co-reviewer, who may not be allowed to review due to ECR status.** These two wrongs do not make a right. How can we remedy this situation? Any policy that requires individuals to simply stop participating in a pervasive practice – and the subsequent policing required to ensure this is resolved – as a way to prevent its undesirable downstream consequences may be ineffective.

We see an analogy to the public health concept of harm reduction, which aims to minimize harm to vulnerable populations while the more challenging work of changing the status quo is tackled, rather than continuing to perpetuate short-term harmful practices because of ideological absolutism (Box 4). Harm reduction interventions as applied to ghostwriting should be “grounded in the recognition that many people are unable or unwilling to stop” (hri.global/what-is-harm-reduction) co-reviewing without due credit – because they see it as a required feature of beneficial peer review training, because they see it as a obligatory status quo and/or necessary delegation of labor in a hypercompetitive environment, because they don’t think to discuss or feel able to discuss authorship, etc. – even when this activity comes at the cost of engaging in widely-accepted unethical behaviors such as not gaining prior permission, breaching manuscript confidentiality policies, or plagiarism.

#### BOX 4: APPLYING PRINCIPLES OF HARM REDUCTION TO GHOSTWRITING

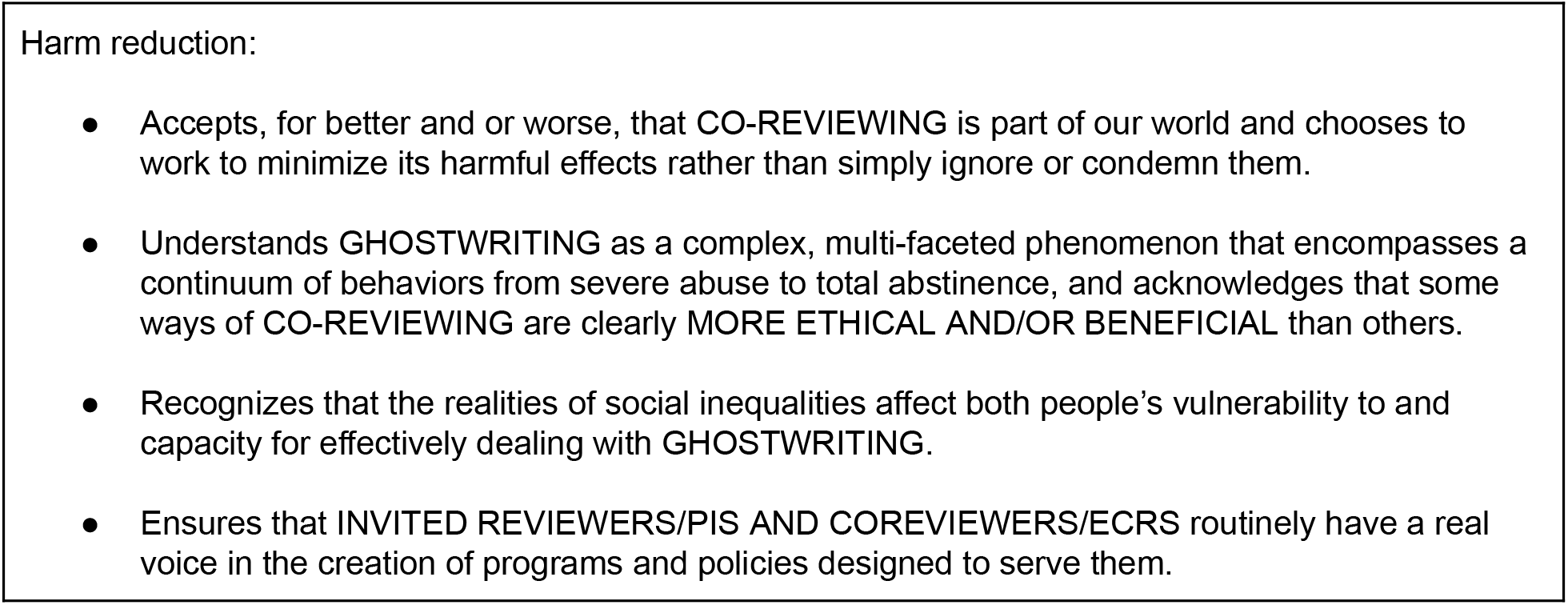
Text in CAPS is original to this manuscript, all other text is quoted verbatim from: harmreduction.orα/about-us/principles-of-harm-reduction/

In summary, we suggest that the best way to reduce the harms of ghostwriting is to 1) make immediate changes to policies that facilitate giving credit where credit is due and ensure that co-review is an effective training exercise and not exploitation of cheap labor while 2) simultaneously working towards promoting long term cultural changes that recognize the role and value gained from involving ECRs in peer review.

## RECOMMENDATIONS

Based on the experiences and opinions of survey respondents presented above, we recommend efforts that encourage ethical practices in quality co-reviewing in peer review. We propose a series of cultural changes for the research community at large (Box A) and practical changes for key stakeholders (Boxes B–D) to improve transparency and the recognition of ECRs in the peer review of manuscripts.

### Cultural changes for the research community at large

#### BOX A: CULTURAL CHANGES FOR THE RESEARCH COMMUNITY AT LARGE

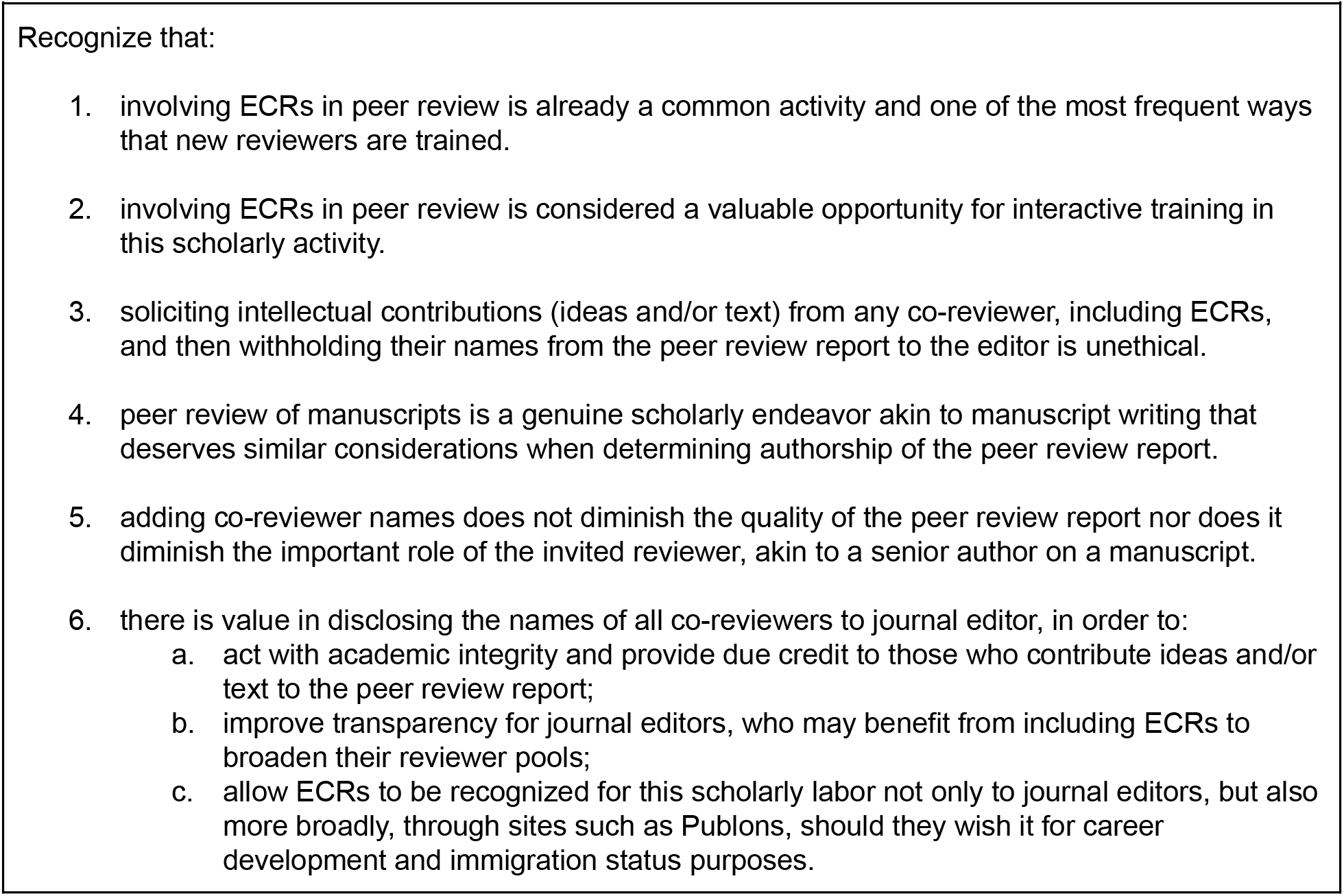

### Practical changes for key stakeholders

#### BOX B: FOR JOURNAL EDITORIAL BOARDS AND COPE

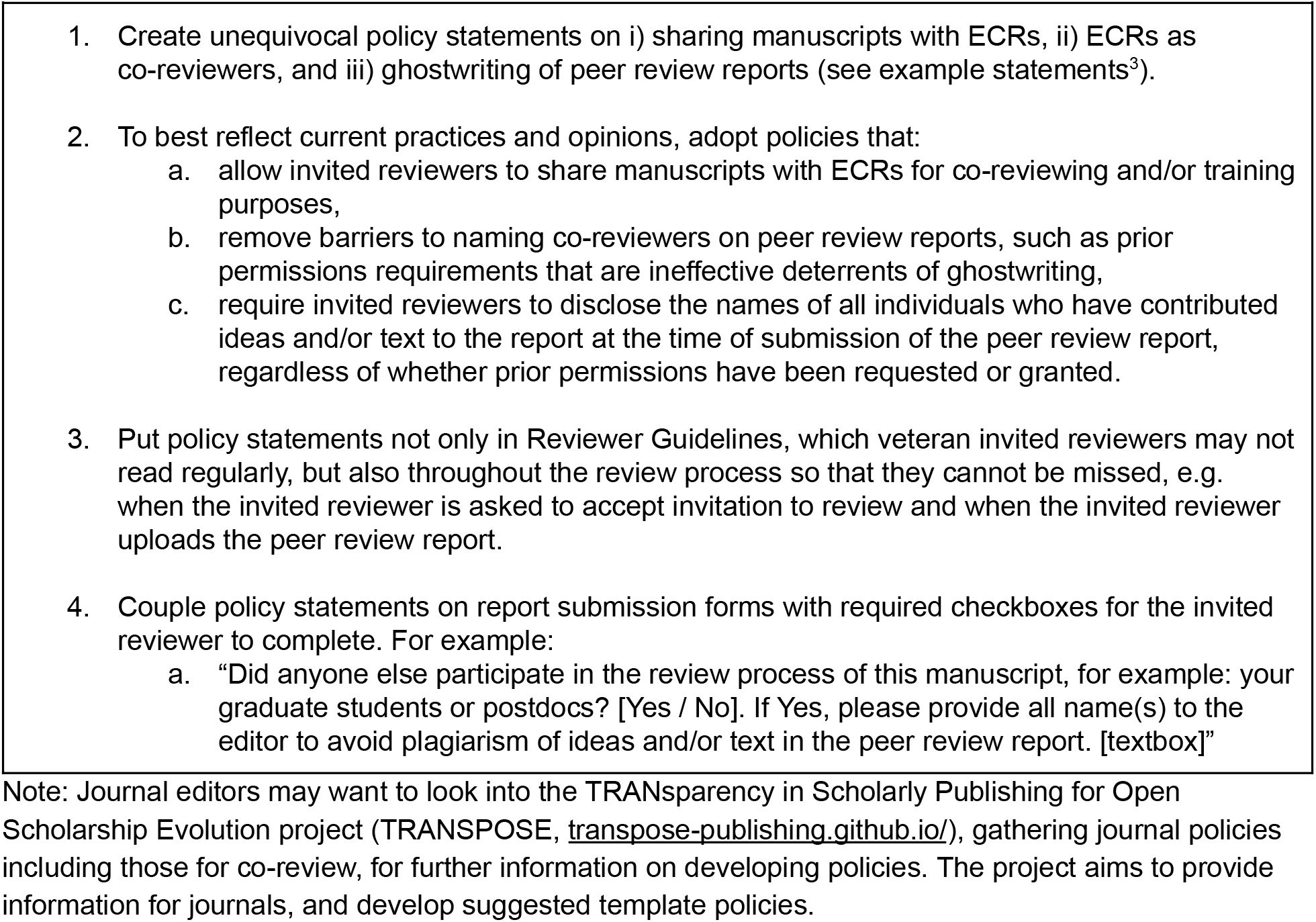

#### BOX C: FOR GRADUATE PROGRAMS, SCIENTIFIC SOCIETIES, ETC.

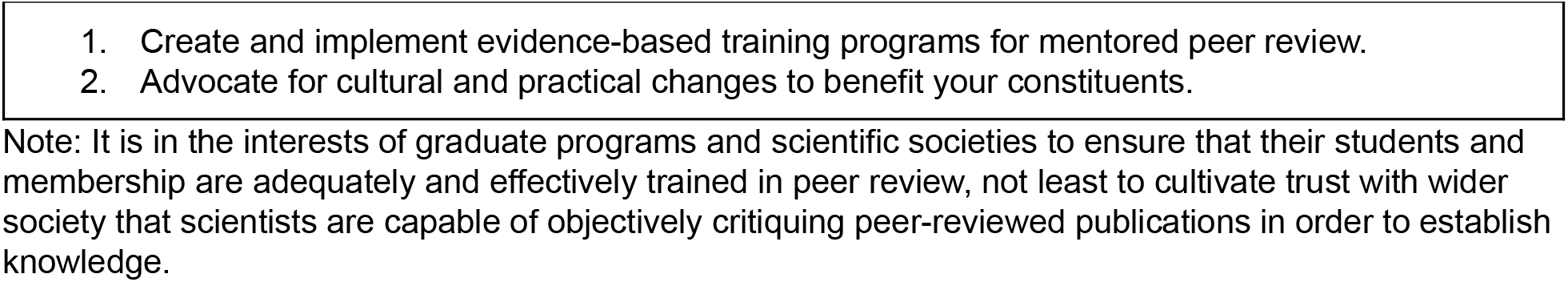

#### BOX D: FOR INVITED REVIEWERS AND COREVIEWERS

**BOX D: FOR INVITED REVIEWERS AND CO-REVIEWERS**
|  |  |
| --- | --- |
| **Both** | Inform yourself of each journal's policies on sharing manuscripts with ECRs, ECR contributions to peer review reports, and ghostwriting. |
|  | Participate in peer review training, especially evidence-based trainings that involve iterative feedback between mentors and mentees. |
| **Invited reviewers** | Inform your trainees of journal policies prior to involving them in peer review. |
|  | Even if journal policies are not clearly stated, initiate a conversation with your trainees about the ethics of co-reviewing, expectations for an ethical division of labor in co-reviewing, and your intentions as to whether you will share ECRs' names to the editor. |
|  | Provide constructive feedback to your trainees on the ideas and/or text that they have contributed to the peer review report, as a training exercise. |
|  | Even if no opportunity is provided to you (e.g. on peer review report submission form), add names of everyone who contributed ideas and/or text to the peer review report on the report itself for the editor to see at the time of submission. |
| **Co-reviewers** | Inform yourself of journal policies prior to agreeing to join an invited reviewer in peer review. |
|  | If comfortable doing so, initiate a conversation with your PI about an ethical division of labor in co-reviewing and your expectations to be/not be named to the journal editor. |
|  | If comfortable doing so, request constructive feedback from your PI on the ideas and/or text that you have contributed to the peer review report, as a training exercise. |
|  | If comfortable doing so, educate others about any value you perceive in being named to the journal editor. |

Note: Responsibility for ethical and effective peer-review training should be a central part of any mentee-mentor relationship, and appropriate training, including feedback from mentors, should be developed in accordance with career stage.

### Differing ability of key stakeholders to affect change

It is important to recognize that the key stakeholders in this process – journals, invited reviewers/PIs, and co-reviewers/ECRs – have differing abilities to affect change based on existing power dynamics in the research and publishing enterprise. While ECRs have an important role to play in asking their mentors for named credit on peer review reports, their vulnerable and relatively disenfranchised role in this hypercompetitive research environment may make them least able and/or willing to advocate for change to the status quo, especially when this can be further affected by concerns arising due to factors such as immigration status. At the next level, invited reviewers in isolated conversations with journal editors may not feel able and/or willing to critique journal policy, especially for journals in which they hope to publish themselves. Journals, therefore, may have the greatest ability to lead on this issue by creating policies that recognize the role that ECRs play in the peer review process, and the value of that role to ECRs themselves, invited reviewers, editors, and the research community at large.

### Benefits of implementing recommendations

All key stakeholders and especially journals stand to benefit significantly from making these common sense changes that recognize ECRs as frequent and valuable contributors to the peer review system. Box E summarizes what we see as the advantages of discouraging ghostwriting and encouraging co-reviewing as training in peer review. We are optimistic that, together, we can change the cultural expectations and policies regarding ECR participation in peer review to reduce harm to vulnerable members of the community and to support the implementation of transparent and ethical practices that align with community experiences and values.

#### BOX E: BENEFITS OF ENCOURAGING CO-REVIEW AS TRAINING AND DISCOURAGING GHOSTWRITING

| **BOX E: BENEFITS OF ENCOURAGING CO-REVIEW AS TRAINING AND DISCOURAGING GHOSTWRITING** |  |
| --- | --- |
| For journals | Increase in pool of potential reviewers* |
|  | Increase in the likelihood that PI/invited reviewer will accept invitation to review when they can do so with co-reviewer help* |
|  | Naming co-reviewers improves transparency, allowing editors to know exactly who contributed to the review |
|  | Co-review as training results in higher quality peer review reports |
| For ECRs, co-reviewers | Named recognition of scholarly labor, with the added potential benefits of:<br>• Being known to the scientific editors<br>• Invitations to review independently<br>• Recognition of peer review experience for career advancement or immigration status |
|  | Become a better peer reviewer through co-review as training |
| For PIs, invited reviewers | Become a better peer reviewer (and mentor) through training co-reviewer |
|  | Involving ECR as co-reviewer decreases workload for overburdened PIs |
| For the peer review system, and so scientific enterprise, on the whole | Ethically better system that recognizes all participants fairly |
|  | Objectively better system with higher quality, less biased peer review |
|  | Improved clarity of expectations for all participants |
|  | Improved integrity of system improves public trust in science |

\* indicates ideas supported by a Reviewer Experience Lead at Elsevier (Mehmani 2019).

## METHODS

### Systematic Literature Review

#### Search procedures

The following procedures were used to perform a systematic review of the peer-reviewed literature for any research on the topic of ECR participation in manuscript peer review. These procedures were developed under the guidance of a professional librarian (author SO) and were based on the Preferred Reporting Items for Systematic Reviews and Meta-Analyses (PRISMA) criteria ((Moher et al. 2009); see Appendix Figure 1). The databases that were searched cover peer-reviewed literature across the life sciences, public policy and social sciences and were comprised of: PubMed, Psychinfo, Web of Science, and PAIS International. These databases were then searched using the following keyword search strategy: (“early career researcher” OR “graduate student” OR “postdoc” OR “fellow” OR “contingent faculty” OR “adjunct” OR “lecturer” OR “instructor” OR “technician” OR “junior scientist” OR “trainee” OR “lab member” OR “research scientist” OR “postdoctoral fellow” OR “research fellow” OR “teaching fellow” OR “junior researcher” OR “mentee”) AND (“peer review” OR “refereeing” OR “invited reviewer” OR “referee” OR “reviewer” OR “co-reviewer” OR “first time reviewers” OR “reviewer training” OR “review partners” OR “contributing author” OR “co-reviewing” OR “reviewing” OR “journal reviewer policy” OR “reviewer guidelines” OR “instructions for reviewers”). These search terms were designed to be broadly inclusive so as to capture any research article with possible relevance to the topic of ECR involvement in manuscript peer review. The resulting collection of 2103 records were imported into the RefWorks 3 bibliographic management database, and duplicate articles were identified and removed using the “Legacy close match” de-duplication filter, resulting in a de-duplicated set of 1952 articles.

#### Relevance screening

Collected articles underwent two rounds of relevance screening. In the initial round, article titles and abstracts were screened independently by two study authors (JG and GM). Both authors used the same inclusion criteria to sort search results into “relevant,” “maybe relevant,” and “not relevant” categories. The criteria for article inclusion were: written in English, published in a peer-reviewed journal, mention of ECRs, and mention of peer reviews of manuscripts. Any article that did not meet the inclusion criteria above was excluded as well as database hits for conference proceedings and dissertations.

118 unique articles remained in the “relevant” and/or “maybe relevant” categories at this stage of screening (see <u>Appendix: Results of relevance screening for literature review</u>). Articles categorized as “relevant” by both initial screeners were selected for full text review (n = 3). Articles categorized as “maybe relevant” by both initial screeners and articles that were differentially categorized as “relevant” vs. “maybe relevant” or “not relevant” by the initial screeners (n = 51) underwent a second round of evaluation by a third, independent screener (author RL) to either confirm categorization as “relevant” or recategorize as “not relevant” to the topic of ECR participation in the peer review of manuscripts. A resulting list of 36 “relevant” articles underwent a full text reading with specific attention being paid to: research question, motivation for article, method of study including details concerning study participants, relevant results and discussions, discussion of peer review and ECRs, and possible motivations for author bias. Of the articles that were found to be “not relevant” to the topic of ECR participation in the peer review of manuscripts for publication in a journal, most discussed other forms of peer review outside the scope of publishing manuscripts (e.g. students engaging in peer review of each other’s written work in a classroom setting as a pedagogical exercise).

### Survey of peer review experiences and attitudes

#### Survey design

We designed a survey to evaluate the peer review experiences of researchers with a specific focus on ghostwriting of peer review reports. The survey was verified by the Mount Holyoke Institutional Review Board as Exempt from human subjects research according to 45CFR46.101 (b)(2): Anonymous Surveys – No Risk on 08/21/2018. All survey respondents provided their informed consent prior to participating in the survey. The survey comprised 16 questions presented to participants in the following fixed order:

- 6 **demographic** questions that collected data on their professional status (current institution, field of research, and career stage) and personal information (gender identity, race/ethnicity, and citizenship status in the United States);
- 7 questions that collected data about their **experience** participating in the peer review of manuscripts for publication in a journal:

- 2 questions about their experience with independent reviewing vs. co-reviewing
- 4 questions about receiving credit for reviewing activities, and
- 1 question about whether and how respondents’ received training in peer review;
- 3 questions that collected data about their **opinions** about co-reviewing and ghostwriting as practices, regardless of whether they had personal experience with these practices:

- 1 question about their degree of agreement on a 5-point Likert scale (*Strongly Disagree; Slightly Disagree; No Opinion; Slightly Agree; Strongly Agree)* with 11 statements about the ethics and value of co-reviewing and ghostwriting,
- 1 question asking their opinion about why ghostwriting as a practice may occur, and
- 1 exploratory future direction question asking if their opinions would change if the names of peer-reviewers were made publically available (“open peer review”). Throughout the survey, there were many opportunities to provide write-in responses in addition to the multiple choice answers. The full text of the survey can be found in the Appendix: Text of The Role of Early Career Researchers in Peer Review – Survey.

#### Survey distribution, limitations, and future directions

The survey was distributed online through channels available to the nonprofit organization Future of Research including via blog posts (McDowell and Lijek 2018), email lists, social media, and word-of-mouth through colleagues. The main survey data collection effort was from August 23 to September 23, 2018. The survey had gathered 498 responses at the time of data analysis.

We recognize that conclusions drawn from any survey data are limited by the size and sample of the population that is captured by the survey. We sought to address this limitation first by collecting as large and geographically and institutionally diverse of a population of ECRs as possible within the month-long timeframe we set for data collection. Secondly, we wished to preemptively address the concern that our survey distribution efforts were inherently biased towards those with strong opinions on the subject and/or those who self-select to receive communication from Future of Research (e.g. listservs, Twitter followers). We therefore attempted to create a “negative control” comparison group of participants who received our survey from channels independent of Future of Research. We created a separate survey form asking identical questions and personally asked 25+ PIs known to the authors, as well as organizational collectives of PIs, to distribute this survey link to their own networks (e.g. lab members, departments). Both surveys were live during the same month-long time period; however, the Pl-distributed survey gathered only 12 responses and so was not sufficient to be used in the analyses presented here. Since the goal of the second, Pl-distributed survey was to be independently distributed outside of our efforts, we are not able to determine whether it garnered so few responses because of a lack of genuine distribution or because the populations it reached were not motivated to participate in the survey. Therefore any conclusions drawn from this study reflect the 498 experiences and perspectives of those individuals so moved to participate in the survey distributed by Future of Research and our results should be considered in this context. One possible future direction for this study is to reopen the survey in conjunction with publication of this manuscript in an effort to broaden and diversify the sampled population, to compare subsequent rounds of responses to our initial 498 responses, and to improve clarity on survey questions (see <u>Appendix: Future directions for survey questions</u>).

#### Survey data analysis

Survey data were analyzed using Microsoft Excel, Version 16 and IBM SPSS Statistics for Macintosh, Version 25 (SPSS^®^ Inc., Chicago, IL, USA). Whenever statistical analyses were used, the exact tests and p values are reported in the appropriate figure legend and/or results text. A p value of less than 0.05 was considered significant. Where a number of demographics are combined in the reporting throughout this study, any analysis group with less than 20 respondents is reported as “<20” instead of the raw n value in an effort to protect the identity of participants.

## Acknowledgements

The authors thank Drs. Jessica Polka, Adriana Bankston, Carrie Niziolek, and Chris Pickett for their constructive comments on this manuscript. GM is grateful to the Open Philanthropy Project which provides support for his role in Future of Research.

## Author disclosures and contributions

Author Gary McDowell is the Executive Director of the 501 (c)3 non-profit Future of Research, Inc. for which he receives financial compensation. No other authors received any source of funding for this work. Rebeccah Lijek formerly served as a volunteer member of the Board of Directors of Future of Research, Inc.

Authors GM and RSL designed the study, wrote and carried out the survey, supervised survey data analysis, supervised the literature review, and wrote the manuscript. JDK performed the data analysis and contributed to the survey design and manuscript. JG and SO performed the systematic literature review and contributed to the manuscript.

## APPENDICES

### Results of relevance screening for literature review

List of all search records categorized by initial screeners as “relevant” and/or “maybe relevant” to the topic of ECR involvement in the peer review of manuscripts based on title and abstract (n = 118). Y = yes, relevant; M = maybe relevant; N = not relevant; – = not evaluated.

| Article title<br>(alphabetical order) | Initial Screeners |  | Tie-breaker | Full Text Reading |
| --- | --- | --- | --- | --- |
|  | #1 | #2 | #3 |  |
| A Fine Balance: How Authors Strategize Around Journal Submission | N | M | - |  |
| A guide to critiquing a research paper. Methodological appraisal of a paper on nurses in abortion care | M | N | - |  |
| A Mentoring Opportunity: A Joint Effort in Writing Letters of Recommendation | N | Y | N |  |
| A model for scholarship in nursing: The case of a private liberal arts college | N | M | - |  |
| A Pragmatic Approach to Getting Published: 35 Tips for Early Career Researchers | M | Y | Y | Y |
| A Survey of Methods for Improving Review Quality | N | M | - |  |
| A web-based synchronous collaborative review tool: A case study of an on-line graduate course | N | M | - |  |
| Advise to research trainees in biomedical sciences - A very personal tribute to the late Dr. Susumu Ohno | N | Y | N |  |
| An ounce of prevention: An associate editor's view | N | M | - |  |
| Assessing postgraduate student perceptions and measures of learning in a peer review feedback process | M | M | N |  |
| Assessment for Learning Research and Writing Skills through Scaffolded Online Peer Review | N | M | - |  |
| Becoming a peer reviewer: engaging in sharing and gaining knowledge | M | Y | Y | Y |
| Being a researcher is not only a matter of publishing: Learning to review scientific articles | Y | Y | Y | Y |
| Biosecurity and the review and publication of dual-use research of concern | N | M | - |  |
| Building Scholarly Writers: Student Perspectives on Peer Review in a Doctoral Writing Seminar | M | M | N |  |
| Calibrated Peer Review: Enabling Technology for a Large - Scale Writing Intensive Neuroscience Course. | M | M | N |  |
| Calibrated Peer Review: Implementation of a computer-based writing-to-learn tool in a large lecture-based introductory biology class | M | M | N |  |
| Calibrated peer review: The application of statistical concepts to a biological research question | N | M | - |  |
| Can Online Peer Review Assignments Replace Essays in Third Year University Courses? And if so, What are the Challenges? | N | M | - |  |
| CAREERS eLife and early career researchers | N | Y | Y | Y |
| Celebrating Peer Review Week | M | M | Y | Y |
| Chinese Early-Career Researchers' Scholarly Communication Attitudes and Behaviours: Changes Observed in Year Two of a Longitudinal Study | M | Y | Y | Y |
| Conducting mental health research: Key steps, practicalities, and issues for the early career researcher | M | N | - |  |
| Developing Knowledge Generation, Communication and Management In Teacher Education: A Successful Attempt of Teaching Novice Computer Users | N | M | - |  |
| Developing laboratory skills by incorporating peer-review and digital badges | N | M | - |  |
| Do peer review models affect clinicians' trust in journals? A survey of junior doctors | M | Y | Y | Y |
| Does mentoring new peer reviewers improve review quality? A randomized trial | Y | Y | Y | Y |
| Does peer review predict the performance of research projects in health sciences? | N | M | - |  |
| Doing peer review and receiving feedback: impact on scientific literacy and writing skills | M | M | Y | Y |
| Doing Peer Review: Reflections From an International Group of Postdoctoral Fellows | M | Y | Y | Y |
| Editorial training models for early-career psychiatrists | N | Y | Y | Y |
| Emergent areas to visualize by the journal strategy holders | M | N | - |  |
| Enhancement of Resident Competencies Via Participation in the Peer Review/Quality Improvement Processes and Morbidity and Mortality Presentations | M | N | - |  |
| Evaluation of multi- and interdisciplinary research - the no-peer problem | N | M | - |  |
| Examining the Effects of Trained Peer Feedback on EFL Students' Writing | N | M | - |  |
| Expanding Group Peer Review: A Proposal for Medical Education Scholarship | M | Y | Y | Y |
| Facilitating improvements in laboratory report writing skills with less grading: a laboratory report peer-review process | N | M | - |  |
| Feedback on Peer Feedback in EFL Composing: Four Stories | N | Y | N |  |
| Graduate student-run course framework for comprehensive professional development | N | M | - |  |
| Guidelines for assessment of publications for contribution to scholarship: a view point. | N | Y | Y | Y |
| Heroes of peer review: Robert Lowe | N | M | - |  |
| How early-career researchers are shaping eLife | N | Y | Y | Y |
| How to achieve accurate peer assessment for high value written assignments in a senior undergraduate course | N | M | - |  |
| Improving peer reviewing in scientific conferences | M | M | Y | Y |
| Increasing the Quality of Peer Feedback in a Professional Writing Course | N | M | - |  |
| Instructors' uses, experiences, thoughts and suggestions regarding Calibrated Peer Review | M | M | N |  |
| Integrating critical analysis, peer review, and independent research in the laboratory phase of a microbiology course | M | M | N |  |
| Investigating peer review as a systemic pedagogy for developing the design knowledge, skills, and dispositions of novice instructional design students | M | N | - |  |
| Judging plagiarism: a problem of morality and convention | N | Y | Y | Y |
| Korean University Students, Attitudes towards Peer Review in EFL Writing | N | M | - |  |
| Learning by reviewing | N | Y | Y | Y |
| Literature search in medical publications | N | M | - |  |
| Manuscript peer review: A helpful checklist for students and novice referees. | M | M | Y | Y |
| Mentor/mentee relationship with the focus on meeting promotion/tenure guidelines | N | M | - |  |
| Misrepresentation and responsibility in medical research | M | M | N |  |
| Mixed Method Study Examines Undergraduate Student Researchers' Knowledge and Perceptions About Scholarly Communication Practices | M | Y | Y | Y |
| Moving from trainee to junior: faculty: a brief guide | M | M | Y | Y |
| Nature neuroscience | N | M | - |  |
| New Reviewer Mentoring Program | N | Y | Y | Y |
| Observations: The Introduction of "Junior Editor" Posts Within Journals | N | M | - |  |
| Open Access Open Grad Students | N | Y | Y | Y |
| Open peer review: a randomised controlled trial | M | M | Y | Y |
| Peer Assessments Using the Moodle Workshop Tool | N | M | - |  |
| Peer review as an educational strategy to improve academic work: An interdisciplinary collaboration between communication disorders and nursing | N | M | - |  |
| Peer Review in Radiology: A Resident and Fellow Perspective | M | N | - |  |
| Peer review process: Guidelines for clinical and forensic psychological reviews | N | M | - |  |
| Peer review: why we need nurse leaders to serve as reviewers for nursing publications | M | M | - |  |
| Peer-Review - using a Paired-Comparison Technique | N | M | - |  |
| Perception of Academic Writing: A Case Study of an ESL Writer during Peer Review | N | M | - |  |
| Perspectives from early career researchers on the publication process in ecology - a response to Statzner & Resh (2010) | M | Y | Y | Y |
| Pit-bull reviewing, the pursuit of perfection and the victims of success | N | M | - |  |
| Plagiarism: an egregious form of misconduct | M | Y | Y | Y |
| Plagiarism. A fools' errand | M | Y | Y | Y |
| Potential of information technology in dental education | N | M | - |  |
| Power of peer review: An online collaborative learning assignment in social psychology | N | M | - |  |
| Preparing the senior or graduating student for graduate research | M | Y | Y | Y |
| PRO myths vs. facts | N | M | - |  |
| Professional Skills Courses Increase Trainee Skills in Writing, Reviewing, Networking, and Poster Presentations | M | M | N |  |
| Professionalism and Communication Education in Pediatric Critical Care Medicine: The Learner Perspective | M | N | - |  |
| Promoting operational research through fellowships: a case study from the South-East Asia Union Office | M | N | - |  |
| Publish or Perish: A Mandate With Negative Collateral Consequences | M | N | - |  |
| Pursuing the journal mission | M | Y | Y | Y |
| Religion, Rebel Scientists, and Peer-Review - 3 Hot Topics | N | M | - |  |
| Replacing the Traditional Graduate Chemistry Literature Seminar with a Chemical Research Literacy Course | M | N | - |  |
| Republished paper: Assuring validity of multisource feedback in a national programme | M | N | - |  |
| Research as a subject--for research | N | M | - |  |
| Research Ethics II: Mentoring, Collaboration, Peer Review, and Data Management and Ownership | N | Y | Y | Y |
| Responsible Conduct of Research in Communication Sciences and Disorders: Faculty and Student Perceptions | M | M | Y | Y |
| Review, revise, and resubmit: The effects of self-critique, peer review, and instructor feedback on student writing | N | M | - |  |
| Reviewing a Manuscript: Disparity Amongst Peer Reviewers' Priorities from Basic Health Sciences and Clinicians | M | Y | Y | Y |
| Student peer review decisions on submitted manuscripts are as stringent as faculty peer reviewers | Y | Y | Y | Y |
| Supersizing e-learning: What a Col survey reveals about teaching presence in a large online class | N | M | - |  |
| Survey of publication outlets in early childhood education: Descriptive data, review processes, and advice to authors | M | N | - |  |
| The "Peer" in "Peer Review" | N | M | - |  |
| The 5th Annual AABT Postdoctoral Panel and Overview: November 2004 | N | M | - |  |
| The Acta Psychiatrica Scandinavica Trainee Advisory Board: Education, mentoring, and experience with the editorial process | M | Y | Y | Y |
| The association between four citation metrics and peer rankings of research influence of Australian researchers in six fields of public health | M | M | N |  |
| The ethics of peer review: What to know before saying "yes" | M | Y | Y | Y |
| The Fox and the Crow' or 'the Foolishness of Vanity Publishing in Fake Academic Journals': A Story from the Arabian Gulf | N | M | - |  |
| The impact of E-learning in medical education | N | M | - |  |
| The invited review ? or, my field, from my standpoint, written by me using only my data and my ideas, and citing only my publications | M | N | - |  |
| The lecturer should know what they are talking about: Student union officers perceptions of teaching-related CPD and implications for their practice | N | M | - |  |
| The More You Know: The Impact of Publication and Peer-Review Experience on Psychology Graduate Students | M | Y | Y | Y |
| The peer review gap: A longitudinal case study of gendered publishing and occupational patterns in a female-rich discipline, Western North America (1974-2016) | M | Y | N |  |
| THE PIPELINE Scientific Teaching in Practice | N | M | - |  |
| The planning and implementation of a faculty peer review teaching project | N | M | - |  |
| The Use of Mock NSF-type Grant Proposals and Blind Peer Review as the Capstone Assignment in Upper-Level Neurobiology and Cell Biology Courses. | M | M | N |  |
| The value of peer feedback opportunities for students in writing intensive classes | N | Y | N |  |
| Three tough acts to follow | N | M | - |  |
| Two Modes of Peer Review among Adult Writers: Which Is Better? | N | M | - |  |
| Use of Peer-Review System for Enhancing Learning of Programming | N | M | - |  |
| Using an Undergraduate Immunology Laboratory Course to Integrate Scholarship and Teaching | N | M | - |  |
| Using Calibrated Peer Review to Teach Basic Research Skills | M | M | N |  |
| Validity and reliability of scaffolded peer assessment of writing from instructor and student perspectives | N | M | - |  |
| Variability in students' evaluating processes in peer assessment with calibrated peer review | M | Y | N |  |
| Where does all the research go? Reflections on supporting trainee-applied psychologists to publish their research | M | Y | N |  |
| Why not assess another trainer and his/her trainee? | N | M | - |  |
| Writing to learn: an evaluation of the calibrated peer review program in two neuroscience courses | M | Y | Y | Y |

**Appendix Figure 1:**
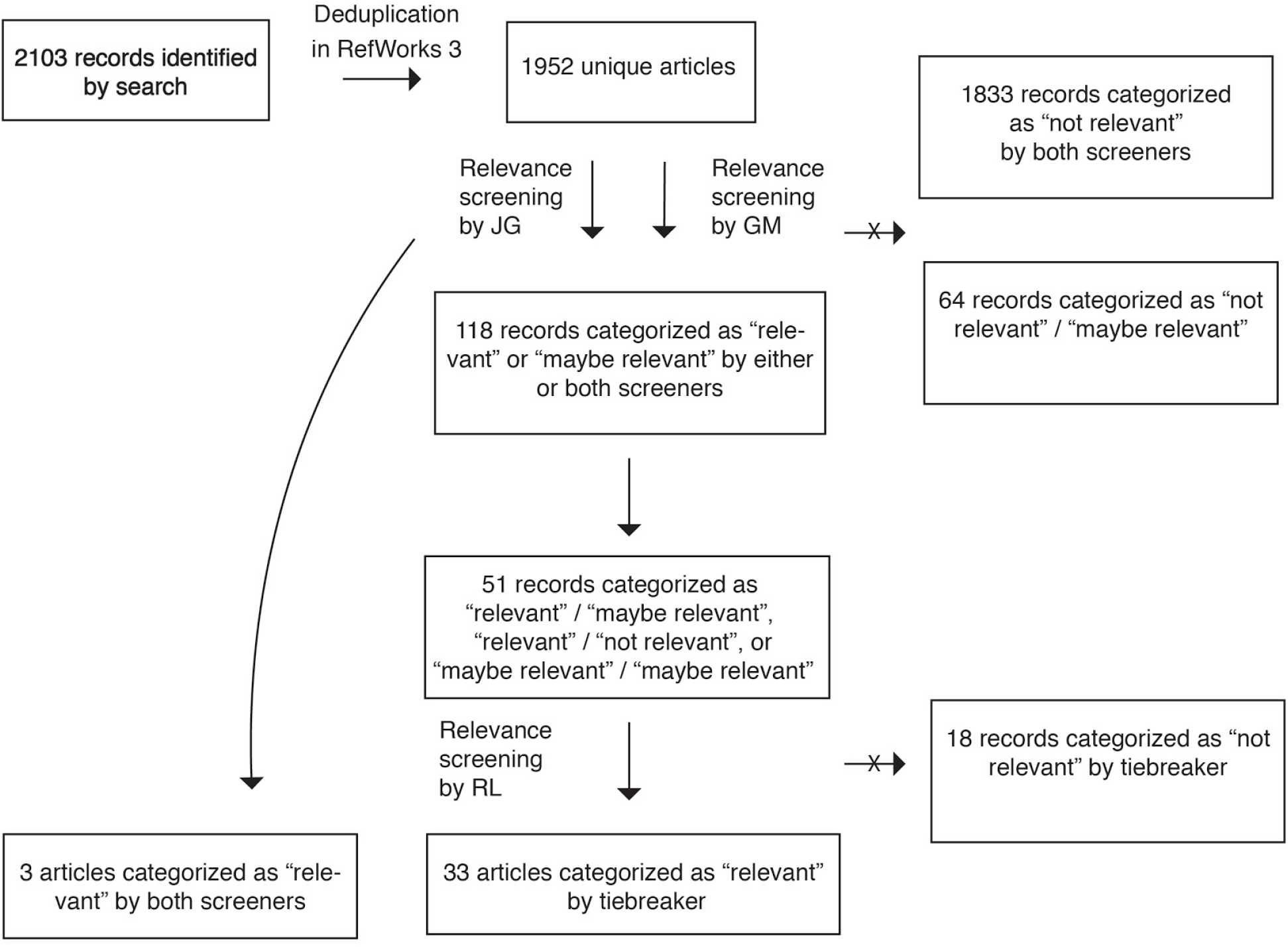
Search strategy for literature review with number of records remaining at each stage.

### Literature on ECR involvement in peer review as a training exercise

Many of the articles uncovered by our systematic literature review on the topic of ECR involvement in peer review of manuscripts that *did not* address our desired topic of ghostwriting instead discussed ECR involvement in peer review of manuscripts as a training exercise for the ECR. Here, we summarize their findings.

#### Peer review training programs with positive outcomes

Trends from the literature indicate several components of a successful peer review training program and where such programs are currently in process. Authors studying the peer review training process for new reviewers tend to conclude that students of peer review learn best by participating themselves in the review process while receiving feedback from more senior reviewers. Successful training programs, in which the reviewers expressed that they had benefited from the training or were evaluated and determined to have benefited from the training, tended to include participation in several rounds of review followed by feedback and revising. Additionally, successful programs often expanded feedback by directing the expert reviewers to report on the same manuscripts as the trainees. This method provided trainees with specific written feedback as well as a pertinent example report to reference. Authors of successful program studies and authors with general policy recommendations for peer review training strategies conclude that a hands-on, iterative process of peer review training with regular and specific feedback are components which positively benefit the peer review trainee (Castelló et al. 2017; Doran et al. 2014).

#### Peer review training programs without positive outcomes

However, one reviewer training study in the pool generated negative results. In their paper, Houry, Green and Callaham found that after a period of training involving mentorship from an expert reviewer to a new reviewer, no differences in mean reviewer quality scores between the mentored and unmentored groups was found. The study goes on to conclude that this similarity in group quality scores is dependent on the mentoring relationship: the expert reviewer mentors were not given any training on how to offer feedback to the trainees. This aspect of the program was deliberately constructed in an attempt to model a training program in which there would be minimal stress on the expert mentors. However, it appears that this ultimately led to an inconsequential training program. Accounting for this information, the authors conclude that a mentorship program should include training and guidelines for mentor-mentee communications which allow for regular feedback from expert to trainee (Houry et al. 2012).

#### Institutions where peer review training takes place

Sources mentioned two primary institutions where training in peer review may take place. Training institutions were identified, with undergraduate education as the first opportunity for training, followed by graduate programs. Journals were the other institutions identified.

The majority of training programs represented in the sources take place within a journal setting. Journal programs tend to include ECRs, such as undergraduate or graduate students, joining the editorial board for a set time period or acting as a reviewer (Castelló et al. 2017; Doran et al. 2014; Houry et al. 2012; Navalta and Lyons 2010; Patterson and Schekman 2018; Picciotto 2018; Harrison 2009).

Training may also take place within undergraduate or graduate institutions. One peer review training study in the found literature took place in an undergraduate setting. Despite the lack of associated literature, Riehle and Hensley determined that undergraduate students are interested in learning about the peer review process (Riehle 2017). In a training study, the Calibrated Peer Review™ system was employed in two undergraduate classes to facilitate students to peer review the work of their classmates while minimizing the extra workload such an exercise might otherwise entail for the professor (Prichard 2005). The participating students were in two separate courses, an introductory neuroscience course and a more advanced neuroscience course. Students in the advanced class did not perform better on peer review exercises than the introductory students, suggesting that until that point, advanced students had not been exposed productively to peer review practice. Authors deemed this to be a successful method for exposing undergraduate students to the peer review process while requiring a realistic time commitment from the course instructor.

While no papers were found detailing the effectiveness of peer review training within graduate institutions, several sources did indicate graduate student perceptions about their program peer review training. In a study including psychology masters and PhD students, a large proportion of participants indicated that their education had lacked in providing information on the peer review and revision process as well as information about how to practice review (Doran et al. 2014). These students indicate that this was a negative aspect of their programs, saying more opportunities for peer review practice should be made available. Authors of this article do indicate that this information may not be generalizable to graduate students as a group because the participating students were found when they pursued a journal review program, something students with adequate peer review education may not be likely to do.

#### Self-facilitated training

Merry et al. provides a list of recommendations for ECRs to facilitate their own training of peer review (Merry et al. 2017). Authors advise working with the mentoring faculty as well as contacting journals directly to seek out peer review opportunities. If mentors are able to give consistent feedback regarding the trainees peer review, it seems that this could be a positive environment in which to learn the skill.

#### Roadblocks to positive outcomes in training programs

Papers discussing journal training programs cite feasibility as the largest roadblock to success (Castelló et al. 2017; Houry et al. 2012). As discussed, it is recommended that peer review training programs for ECRs feature a system which provides regular, specific feedback from expert reviewers. Such programs require high levels of labor, involving organization and time commitment from program leaders and expert reviewers. This is a significant investment for a business which may be in conflict with the desire to maximize journal profit. One possible solution presented involves student-run journals hiring increased numbers of student reviewers and editors so experience may be gained in the field (Doran et al. 2014; Patterson and Schekman 2018). However, this solution does not address the recommendation for expert reviewers to provide feedback.

### Text of The Role of Early Career Researchers in Peer Review – Survey

#### Background

Peer review of academic manuscripts is essential to maintain integrity in science and is integral to the journal publication process. Early Career Researchers (ECRs) often contribute to this peer review process. While ECRs may review manuscripts jointly with or under the direction of a senior academic, such as a Principal Investigator (PI), Group Leader, or Professor, a large number of ECRs claimed in a recent survey to have acted as peer review “ghostwriters”; that is, the peer review report (i.e. the final review submitted to the journal editor) had only the senior academic’s name attributed to the report. For the rest of this survey, we refer to the senior academic as the “PI” and any junior academics under their supervision as “ECRs.”

This survey is designed to collect more data about the phenomenon of ghostwriting by ECRs. The goal of this survey is to assess the experiences and opinions of the community, and to recommend best practices for recognizing co-reviewing activities.

This survey contains 16 questions and is estimated to take 15 minutes.

#### Statements of Disclosure, Ethics and Informed Consent

This survey was created by researchers affiliated with the Future of Research, a non-profit organization in the United States that is promoting an effort to increase transparency about co-reviewing activities by ECRs. You can find out more about our work on ECRs in peer review at our website here: http://futureofresearch.org/ecrpeerreview/

The researchers respect the confidentiality and anonymity of all respondents. No identifiable private information will be collected by this survey. Your participation is voluntary and you can choose to stop at any time. Please complete this survey only once. By choosing to submit answers to this survey, you thereby provide your informed consent to voluntarily share your experiences and opinions with the researchers, who intend to publish a summary of the results of the survey but not the raw data with participants’ individual demographic information.

You may contact Gary McDowell, Executive Director of the Future of Research, at at any time during the study if you have questions or concerns about your participation.

This survey has been verified by the Mount Holyoke Institutional Review Board as Exempt according to 45CFR46.101 (b)(2): Anonymous Surveys – No Risk on 08/21/2018.

I provide my informed consent to participate in this survey.

- Yes

#### Professional Information

Q1.

1. What is your current institution? Fill in blank box (e.g. Harvard School of Medicine).

Q2.

What is your current field of research? Fill in blank box (e.g. biomedicine; physics; philosophy; economics; etc.).

Q3.

What is your current career stage?

- Undergraduate Student
- Graduate Student – Masters
- Graduate Student – PhD
- Postdoctoral Researcher
- Staff Scientist
- Adjunct Professor
- Principal Investigator (PI)
- Other (please describe)

#### Demographic Information

Please feel free to skip any of the following questions if you feel they would be sufficient to uniquely identify you.

Q4.

What is your gender identity?

- Female
- Male
- Prefer not to say
- Other (please describe)

Q5.

What is your race/ethnicity? Select all that apply.

- Asian
- Black or African American
- Hispanic or Latinx
- Native American or Alaska Native
- Native Hawaiian or Pacific Islander
- White
- Prefer not to say
- Other (please describe)

Q6.

If you are based in the U.S., are you a U.S. Citizen/Permanent Resident?

- Yes
- No
- Not based in the U.S

#### Your peer review experience

Q7.

How many times in your career have you reviewed an article for publication independently, i.e. carried out the full review and been identified to the editorial staff as the sole reviewer?

- 0
- 1-5
- 6-20
- 21 +

Q8.

How many times in your career have you contributed ideas and/or text to peer review reports where you are not the invited reviewer (e.g. the invited reviewer is the PI for whom you work)?

- 0 – skip to question 13
- 1-5 – go to question 9
- 6-20 – go to question 9
- 21+ – go to question 9

Q9.

When you were not the invited reviewer, what was the extent of your involvement in the peer review process? Please select all that apply to your entire peer review experience (e.g. across multiple manuscripts).

- I read the manuscript, shared short comments with my PI, and was no longer involved
- I read the manuscript, wrote a full report for my PI, and was no longer involved
- I read the manuscript, wrote the report, my PI edited the report and we submitted the report together with both of our names provided to the editorial staff
- I read the manuscript, wrote the report, my PI edited the report and my PI submitted report with only their name provided to the editorial staff
- I read the manuscript, wrote the report, and submitted it independently without my PI’s name provided to the editorial staff

Q10.

To your knowledge, did your PI ever submit your reviews without editing your work?

- Yes
- No
- Don’t know

Q11.

To your knowledge, did your PI ever withhold your name from the editorial staff when you served as the reviewer or coreviewer?

- Yes – proceed to question 12
- No – skip to question 13
- Don’t know – proceed to question 12

Q12.

Consider cases where you contributed to a peer review report and you know your name was NOT provided to the editorial staff. When discussing this with your PI, what reason did they give to exclude you as a co-reviewer?

- Did not discuss with my PI
- Journal does not allow ECRs to review
- Journal requires prior approval to share manuscript, which was not obtained
- Intellectual contribution not deemed sufficient
- Co-authorship is for papers, not for peer review reports
- Other (please describe)

Q13.

How did you gain training in how to peer review a manuscript? Select all that apply.

- Online resource
- Your PI
- A postdoc in the lab
- A graduate student in the lab
- Journal Club
- Attending an in-person course/workshop
- From receiving reviews on my own papers
- I have had no training
- Other (please describe)

#### Your opinions on peer review

The following questions are about your opinions, not necessarily your experiences. Please answer the following questions regardless of whether or not you have participated in peer review.

A ghostwriter is defined as a person that writes text or other scholarly works without receiving authorship.

These questions are about submitting names of co-authors to the editorial office, not making the identities of reviewers publicly available.

Q14.

Please indicate how strongly you agree with the following statements. You may also submit comments to expand and/or clarify your opinions in the textbox below.

*Options: Strongly Disagree; Slightly Disagree; No Opinion; Slightly Agree; Strongly Agree*

- Involving members of a research group in peer review is a beneficial training exercise.
- It is ethical for the invited reviewer (e.g. PI) to involve others (e.g. their trainees) in reviewing manuscripts.
- It is ethical for the invited reviewer (e.g. PI) to submit a peer review report to an editor without providing the names of all individuals who have contributed ideas and/or text to the report.
- Ghostwriting a peer-review report for your PI is an ethically sound scientific practice.
- When a journal invites a PI to review, that is equivalent to the journal inviting anyone in that PI’s research group with relevant expertise to contribute to the review.
- It would be valuable to have my name added to a peer review report (e.g. to be recognized as a co-reviewer by the editor; or to use a service such as Publons to be assigned credit).
- Anyone that contributes ideas and/or text to the review report should be included as a co-author on the review.
- The only person who should be named on a peer review report is the invited reviewer, regardless of who carried out the review.
- Adding names of other contributors to a peer review report diminishes the credibility of the report.
- The current system used to name and order co-authors on manuscripts in my field should also be used to the name and order co-reviewers on peer review reports.
- The current system used to identify author contributions on manuscripts in my field (e.g. AB did the experiments, AB and CD analyzed the data and wrote the paper), or the CRediT taxonomy (https://casrai.org/credit/), should also be used to identify author contributions on peer review reports (e.g. AB reviewed the experiments, AB and CD wrote the report).

Please submit any extra thoughts or comments regarding question 14 here:

Q15.

What do you think are the reasons why the names of co-authors on peer review reports may not be provided to the editorial staff? Please select all that you think apply.

- A lack of a mechanism (such as a textbox, with language demonstrating expectations that co-reviewers be listed) to include this information in the peer review report submission process.
- A belief that reviews should only be done by the invited reviewer, and not by, or with assistance from, anyone else.
- A belief that only the invited reviewer deserves authorship, even when others contribute ideas and/or text to the review report.
- A belief that there is no strong ethical reason to add co-authors names.
- A belief that including co-author information would demonstrate that the PI breached the confidentiality of the manuscript.
- Some ECRs may not be comfortable asking for co-authorship.
- A belief that keeping ECR names off of peer review protects ECRs during a vulnerable time in their career.
- Some co-reviewers want to be able to write critical reviews anonymously.
- A belief that ghostwriting does not occur: everyone always provides the names of all contributing authors to the editorial office.
- A belief that ghostwriting does not occur: PIs are the only people that contribute to peer review reports.
- Other (please describe)

Q16.

16. Would any of the opinions you have just expressed change if the content of peer review reports (i.e. the text of reviews) were published openly alongside the papers? And should such published reports include or exclude the reviewer’s name(s)? Please explain.

#### Thank you!

You have completed the survey. Many thanks for your responses! Please check http://futureofresearch.org/ecrpeerreview/ or subscribe to our blog to keep updated on the results. Please share the link to the survey with your colleagues: https://tinyurl.com/ECRs-in-peer-review

#### List of institutions with multiple survey respondents

**In alphabetical order:**

|  |  |
| --- | --- |
| Albert Einstein College of Medicine | St Jude Children's Research Hospital |
| Baylor College of Medicine | Stanford |
| Boston Children's hospital | Tufts University |
| Boston University | University of California Berkeley |
| Broad Institute | University of California Los Angeles |
| California Institute of Technology | University of California San Francisco |
| Carnegie Mellon University | Université de Montréal/University of Montreal |
| Cold Spring Harbor Laboratory | University of Alabama at Birmingham |
| Columbia University | University of British Columbia |
| City University of New York | University of Cambridge |
| Dana-Farber Cancer Institute | University of Chicago |
| Dartmouth College | University of Georgia |
| European Molecular Biology Laboratory | University of Illinois at Chicago |
| Fred Hutchinson Cancer Research Center | University of Kentucky |
| Harvard Medical School | University of Massachusetts |
| Harvard University | University of Michigan |
| Icahn School of Medicine at Mount Sinai | University of Minnesota |
| Imperial College London | University of North Carolina at Chapel Hill |
| Indian Institute of Technology Madras | University of Oxford |
| Lawrence Berkeley National Laboratory | University of Pennsylvania |
| Ludwig Maximilian University of Munich | University of Pittsburgh |
| Max Planck Institute | University of Puerto Rico |
| Mayo Clinic | University of Technology of Vienna |
| Memorial Sloan Kettering | University of Toronto |
| New York University | University of Vermont |
| Not Disclosed | University of Washington |
| Ohio State University | University of Wisconsin |
| Okinawa Institute for Science and Technology | University of Würzburg |
| Princeton University | Vienna University of Technology |
| Rockefeller University | Washington University in St. Louis |
| South Dakota State University | Yale University |

#### List of topics assigned to fields of study

**Life Sciences:**

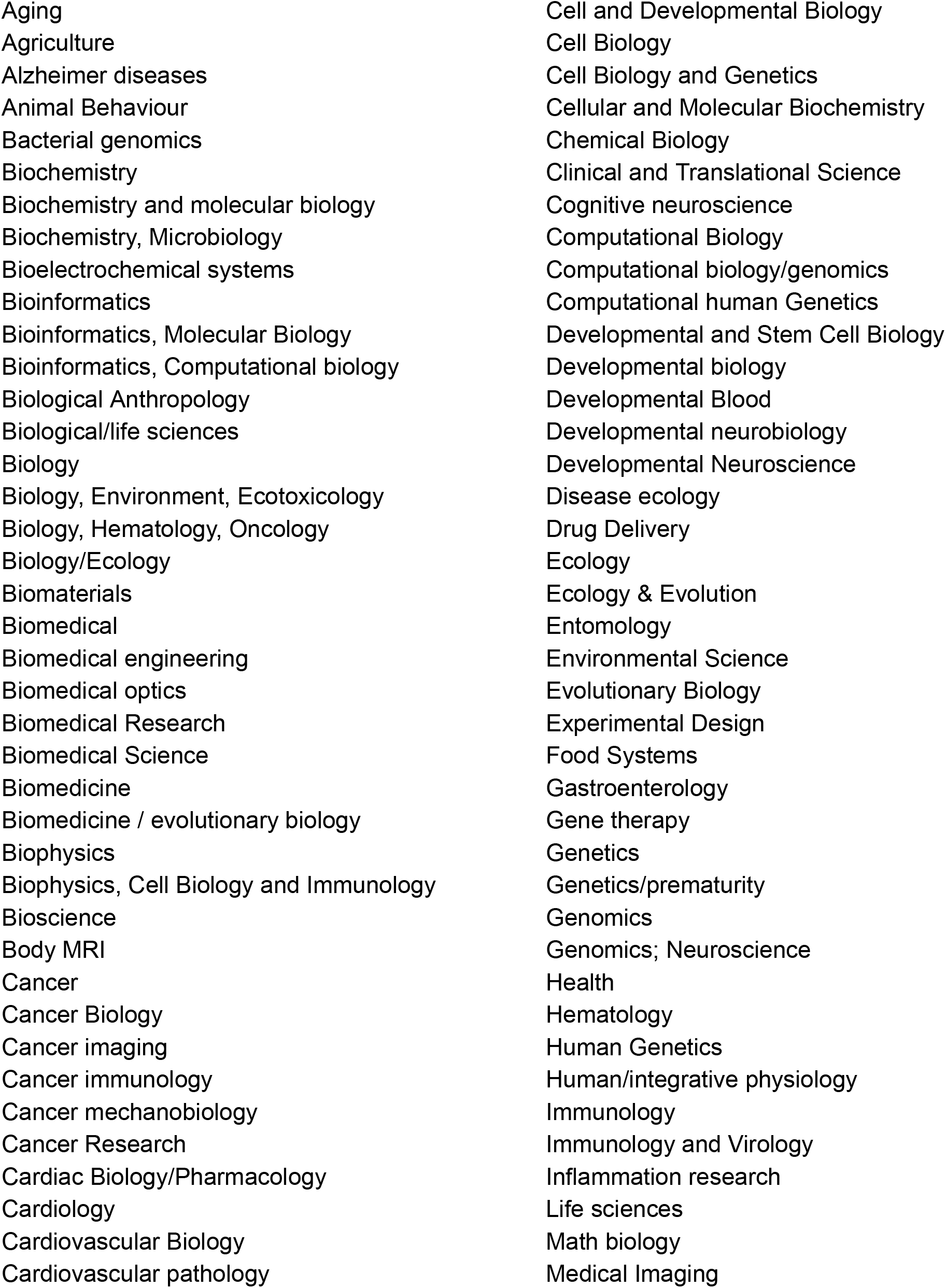
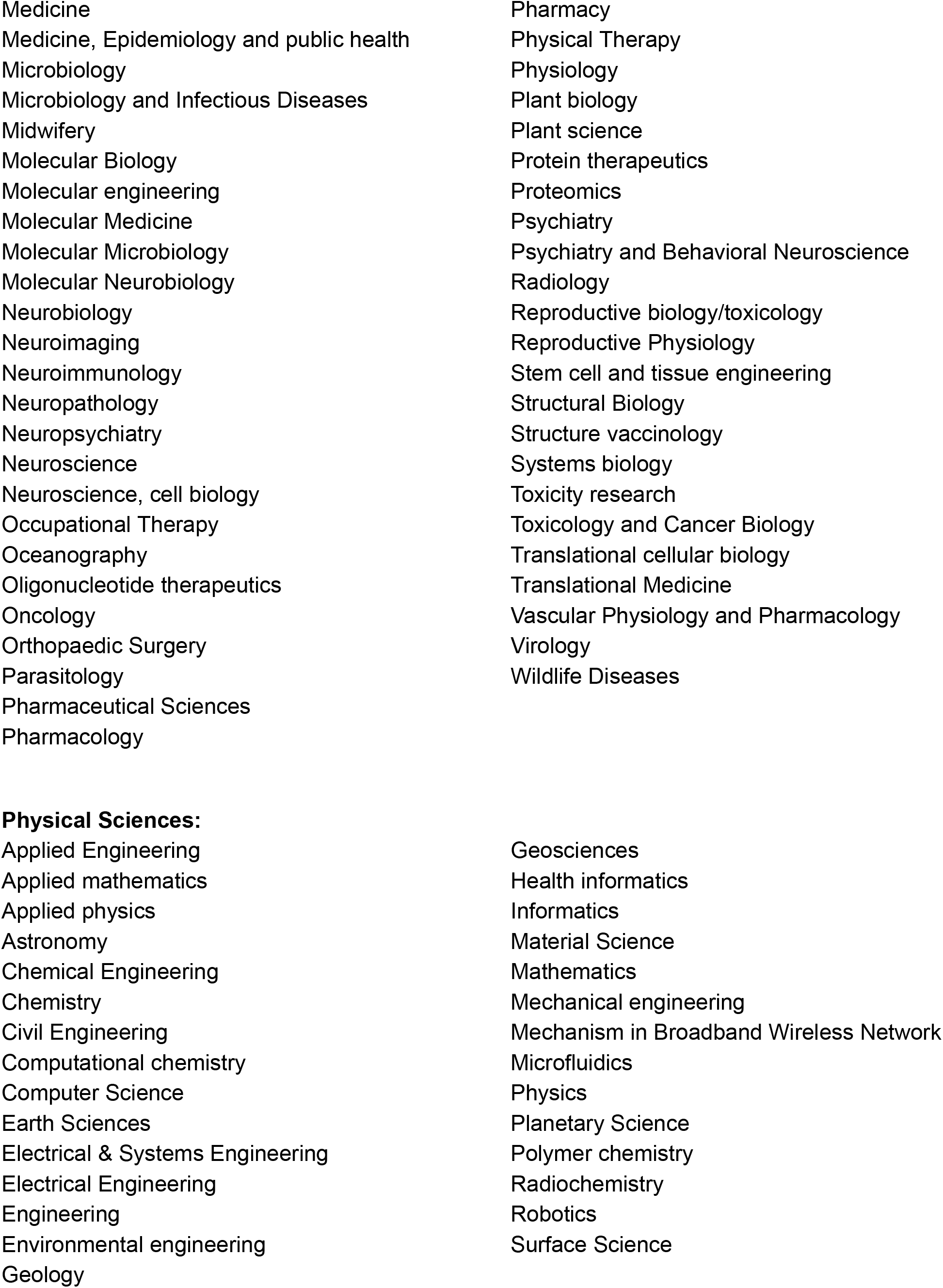
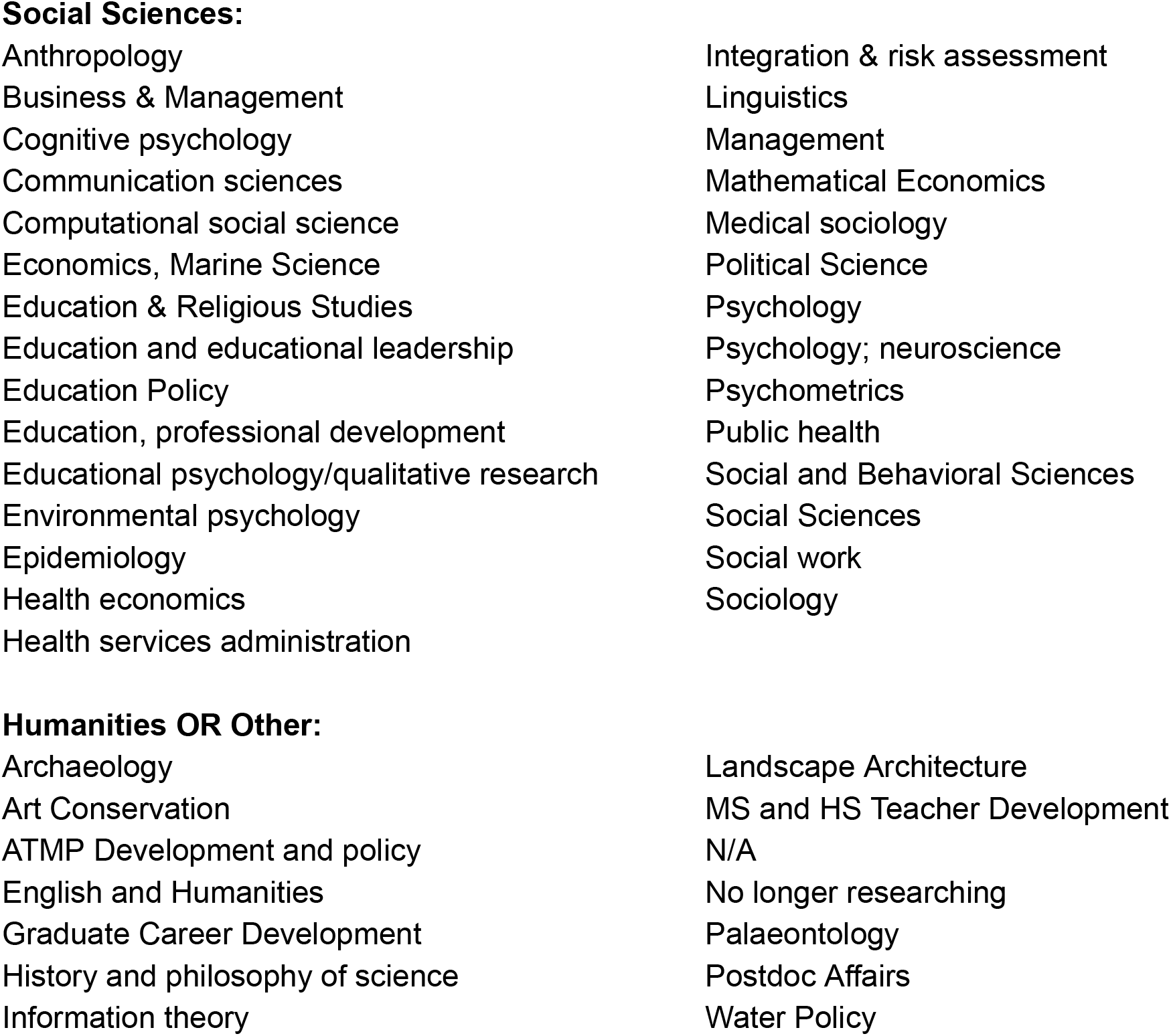

#### Written responses giving reasons PIs excluded survey respondents as co-reviewers

<u>In response to Question 12. Responses are not edited unless noted</u> *[in this fashion]:*

- Apparently this duty is part of my job description
- At my request; wished to remain “anonymous” to manuscript authors
- Didn’t think about including me, didn’t know how to do so
- Good training/practice and could put journal reviewer on my CV ***[subsequent personal information about respondent’s career stage removed]***
- He said only he would be invited to review for such a prestigious journal and “we” need this for future submissions.
- He wanted me to learn under his supervision first. I did this twice, then the third time I reminded him to suggest my name to the editorial staff so that I could get the invite for review myself, which I did receive.
- He was in a hurry and he couldn’t figure out the journal’s website.
- I did not ever submit paper
- I did volunteer to review manuscripts knowing my name was not gonna be listed as a reviewer
- I was told this is how one gets to train to review papers and grants
- In my case I assisted other ECRs with their reviews, and there was no mechanism to indicate multiple reviewers collaborated on the review
- In my opinion my intellectual contribution was not sufficient. It was a training exercise and no one really gets career benefit for reviews anyway. Frankly this is a leading serries of questions.
- It was good for my career to practice.
- Its good experience for me
- Journal doesn’t have co-reviewers.
- no clear answer given
- no reason given other than to help me learn to write reviews
- PI is not fit to sit in editor or reviewers position so only he requests me to review the manuscripts.
- PI signs review and review was v contentious
- Pi surprised I would be interested in being acknowledged, and seemed like too much trouble to acknowledge my contribution. There was no box in the online form to declare it.
- reviewing papers as ECR is part of the ECR training
- The interaction was teh form of a discussion over elements of the review, neither of us considered my contribution as “co-reviewing”
- There wasn’t a field to add my name/ it wasn’t clear if co-reviewers were allowed (I.e. he didn’t know whether it was allowed or not)
- Theres not an option in most cases to list multiple people as reviewers from the same lab, you can still list on CV.
- They forgot
- this is standard practice in ***[place name redacted]-*** it’s never questioned
- This was not explicitly discussed, but the PI implied this is “common practice” and normal for ECRs to gain experience
- tried to discussed but it was ignored by my PI.
- We discussed it but they did not give me a reason
- When I reviewed papers with my PI it was more so an exercise for me, not a situation where my PI did not actually review the work in question and only took my review. It was reviewed jointly as part of a learning/training process for me. I do not know whether or not my comments ended up being incorporated into his final review of the manuscripts.

#### Examples of formalized training in peer review

Some graduate programs, scientific societies, and journals already provide materials for training in peer review. For example, the American Chemical Society’s Publications branch provides the free ACS Peer Review Lab™ course, designed by editors, researchers and publication staff (<u>acsreviewerlab.org</u>). GENETICS, a journal of the Genetics Society of America, provides a peer review training program with virtual training sessions for ECR members (<u>genetics-gsa.org/careers/training_program.shtml</u>). Nature provides an online course on peer review (https://masterclasses.nature.com/courses/205). The Journal of Young Investigators (jyi.org provides training at the undergraduate level. Publons (which provides a mechanism for recording and crediting peer review activity) has a peer review academy online (https://publons.com/community/academy/).

In terms of graduate training, some examples exist at the level of the graduate program. For example, Dr. Needhi Bhalla, at the University of California Santa Cruz, has shared their system (see below for <u>Template from Dr. Needhi Bhalla (UCSC) for peer review training using preprints</u>, adapted from (Halbisen and Ralston 2017)) whereby students review pre-prints in the graduate cell biology class, using a template, and then send reviews to the preprint authors, thereby combining peer review training in journal clubs with the ability to actually make comments on preprints that authors can then use in preparation of a final manuscript for publication at a peer-review journal, should they wish (Avasthi et al. 2018). Examples of other classes that we have so far found or received information on include Class 230 in the PhD Program in Biological and Biomedical Sciences at Harvard; a class on critiquing papers at University of California San Diego; a class in the graduate program in Systems Biology at Harvard Medical School; a requirement to review a paper as a final exam recently introduced at the Graduate Field of Biochemistry, Molecular, and Cell Biology at Cornell University and the University of Texas Southwestern is experimenting with an advanced course combining peer review, literature review, debate and commentary communication and lay-audience oriented writing.

### Template from Dr. Needhi Bhalla (UCSC) for peer review training using preprints

#### Assignment description

“Your assignment is to pick a cell biology preprint from biorxiv (http://biorxiv.org/collection/cell-biology) and review it. This assignment is due [DATE] [TIME], Please submit your review as a word document so that I can edit it.

I’ll assess, edit and grade your review. Afterwards, you will email your edited review to the corresponding author(s), cc’ing me on this email. Your grade is contingent upon submission of your review to the authors.

I’d like you to organize your review as follows:

Part 1. Summary (less than 500 words):

Write a brief overview of the author’s findings and provide a general assessment on the quality of the work: strengths and weaknesses.

Part 2. Detailed comments:

Address each of the questions below, providing specific examples to justify your comments.

1. Significance Does the author provide justification for why the study is novel and how their results will influence the field?
2. Observation Are the author’s descriptions of the data accurate and are all key experiments and hypotheses covered? Are the author’s arguments logically and coherently made? Are counterbalancing viewpoints acknowledged and discussed? Are they sufficiently detailed for a non-expert to follow? Do they include superfluous detail?
3. Interpretation Are the inferences supported by the observations? Do you agree? If not, what experiments would you need to see to be convinced? Please limit any requests for new work, such as experiments, analyses, or data collection, to situations where the new data are essential to support the major conclusions. Any requests for new work must fall within the scope of the current submission and the technical expertise of the authors.
4. Clarity Is the manuscript easy to read and free of jargon, typos, and grammatical or conceptual errors? Is the information provided in figures, figure legends, boxes and tables clear and accurate? Is the article accessible to the non-specialist? Tips: It is important to provide a helpful review that you would want to receive. Critical thinking does not need to be negative to be convincing! Let me know if you’d like to consult with me about your choice of papers or have any questions.”

#### Future directions for survey questions

### Opinions on open peer review reports and public naming of reviewers

The final (16th) question of the survey provided respondents with an open comment box in which to reflect on whether any of their opinions would change if either the *contents* of peer review reports or the *names of reviewers were* openly published to journal readership. This question is a stark contrast to the the rest of the survey questions, all of which instead only ask about naming co-reviewers to the journal staff and editors but not openly to the public readership. We included this question as a test balloon for future surveys that might focus on open peer review vs. traditional models of closed peer review. Due to its tangential relationship to the goals of the current study on ghostwriting and due to the question’s open-ended, write-in nature, we did not perform a systematic analysis of responses as with the other questions. A qualitative summary of responses is below.

61% of respondents chose to write a response to this question, and of these approximately one third reported that no, their opinions would not change if the peer review reports nor names of reviewers were openly shared with the public readership (it should be noted that these comments mostly, but not necessarily, endorsed the specific open peer review features suggested in the question). The remaining respondents expressed a variety of concerns, mostly surrounding the loss of anonymity of reviewers rather than what appeared to be the less controversial concept of publishing the contents of peer review reports. Respondents’ hesitations about anonymity often centered on the effect that naming ECRs and URMs might have on these vulnerable populations. These responses reflect other conversations about open peer review (Polka et al. 2018; Tennant et al. 2017; Ross-Hellauer 2017; Ross-Hellauer et al. 2017) and in the context of recent data about referee behavior in open peer review (Bravo et al. 2019) warrant further analysis beyond the scope of this manuscript.

### Improving clarity in survey questions

One survey question that would benefit from disambiguation in future iterations of the survey is: “Agree/Disagree: When a journal invites a PI to review, that is equivalent to the journal inviting anyone in that PI’s research group with relevant expertise to contribute to the review.” It was brought to our attention by respondents’ emails and write-in comments during the survey response period that there was confusion about this statement. We had hypothesized that we might find agreement with the statement; however, we found a substantial amount of disagreement which may be due in part to the various ways the question may have been interpreted. Our intention was to determine if respondents agreed that, in practice, it could be reasonable for all engaged in journal publication and peer review to expect that an invited reviewer would have various motivations to share a manuscript with the relevant expertise in their research group, particularly in cases where a postdoc is likely more familiar with the literature or experimental techniques than a Principal Investigator overseeing a number of projects. However it became apparent that this question was quite open to various interpretations as described in write-in comments received on the question, including:

- Does the respondent believe this *should* be the case?
- Does the respondent believe this is what is *actually happening*?
- Does the respondent consider that journals have this intent when they invite reviewers?

This therefore renders interpretation of responses to this question difficult, and so we chose not to draw conclusions from this particular result. We aim to clarify this question should there be future iterations of this survey.

Another survey question that would benefit from future disambiguation draws from respondents’ ability to select multiple responses for the question “When you were not the invited reviewer, what was the extent of your involvement in the peer review process? Please select all that apply to your entire peer review experience (e.g. across multiple manuscripts).” and indeed the inability to discern whether respondents supplying only one answer were selecting that response because that comprised the totality of their experiences, or because they selected the most common experience. In comparing responses to this question with other questions, it may be that there are analyses that are affected by the assumption that it is possible to apply one response of many to a response to another question. We attempted to preemptively disambiguate responses by asking whether respondents had “ever” experienced certain things in subsequent questions.

We are also keen to solicit suggestions on possible questions to include or expand upon in a revised study, and possibly removing certain questions. Of course, we also recognize that sharing the results of the first 498 respondents could itself bias subsequent data collections efforts, and we are currently considering which direction to take, particularly as to which questions could potentially be prejudiced by awareness of prior data.

## Footnotes

1 Our search did not recover the survey shared on the INSIDE eLife blog (2018), mentioned above, because it was not published in the peer-reviewed literature nor did it specifically address ghostwriting. Other grey literature would similarly not be included in this dataset.

2 Many journals adhere to The Committee on Publication Ethics (COPE) Ethical Guidelines for Peer Reviewers (https://publicationethics.org/files/Ethical_Guidelines_For_Peer_Reviewers_2.pdf) which state that, “Supervisors who wish to involve their students or junior researchers in peer review must request permission from the editor and abide by the editor’s decision.”

